# Coding of social odors in the hippocampal CA2 region as a substrate for social memory

**DOI:** 10.1101/2021.09.02.458744

**Authors:** Sami I. Hassan, Shivani Bigler, Steven A. Siegelbaum

## Abstract

The ability to encode and update information about individuals is critical for lasting social relationships. Although the hippocampus is important for social recognition memory, its underlying neural representations remain elusive. Here we investigate the neural codes mediating social recognition and learning by examining social odor recognition and associative odor-reward learning in mice. We performed high-resolution calcium imaging from the hippocampal CA2 region of awake head-fixed mice, as CA2 is necessary for social recognition memory. We find that CA2 encodes specific neural representations of novel social odors that are further refined during associative odor-reward learning. Optogenetic silencing of CA2 impairs the formation of reward associations. Furthermore, CA2 population activity represents odors in a geometry that enables abstract representations of social versus non-social odors. Thus, CA2 distinguishes multiple forms of olfactory stimuli to enhance the learning of social odors and associations, which are poised to serve as substrates of social memory.

## Introduction

The encoding and recall of social memory is a challenging problem as it requires that neural circuits encode distinct representations of individual conspecifics and associate those individuals with specific social experiences. This requires the integration of multiple sensory modalities with the outcomes and context of social encounters to represent the dynamic nature of social interactions and allow for successful adaptive behavior. Since the early studies of Scoville and Milner on patient H.M. (Scoville and Milner, 1957), it has been clear that the formation of such complex experience-dependent social representations requires the participation of the hippocampus (Eichenbaum, 2017). Subsequent studies across a range of species have confirmed the importance of the hippocampus in social cognition (Kogan et al., 2000; Sanders and Warrington, 1971; Sliwa et al., 2016; Uekita and Okanoya, 2011). However, our knowledge of the nature and localization of the neural representations for individual conspecifics and how those representations may be refined through experience remains poorly understood.

One relatively recent advance is the finding that a hippocampal subcircuit involving communication from the dorsal region of CA2 (dCA2) to the ventral region of CA1 (vCA1) is central to social memory storage. Thus, silencing or lesioning of dCA2 (Hitti and Siegelbaum, 2014; Oliva et al., 2020; Stevenson and Caldwell, 2014), silencing its inputs to vCA1 (Meira et al., 2018), or silencing vCA1 itself (Okuyama et al., 2016) are all able to prevent social memory formation. Moreover, populations of neurons in these regions form socially relevant representations. Recordings from freely moving mice reveal that pyramidal neuron firing in dorsal CA2 encodes social novelty and can distinguish a novel animal from a familiar littermate (Donegan et al., 2020; Oliva et al., 2020). In contrast, a fraction of neurons in ventral CA1 selectively fire in response to a familiar but not novel animal (Okuyama et al., 2016). What is the nature of the sensory input to hippocampus that enables social discrimination? What are the dynamics by which novelty responses are transformed to representations of social identity? To what extent can social representations be modified by experience, such as associative learning?

To investigate the nature of social representations in CA2, we focused on responses to volatile social odors (urine) from individual mice, as such odors are the most salient sensory signal underlying social memory (Hurst et al., 2001) and other social behaviors (Brennan and Kendrick, 2006) in rodents. As previous studies of social representations in CA1 and CA2 of freely moving mice are complicated by firing responses to spatial location of hippocampal place cells (Alexander et al., 2016; Donegan et al., 2020; Okuyama et al., 2016; Oliva et al., 2020), which are found in both areas, we examined social odor responses using high resolution, large-field two-photon calcium imaging in the CA2 region of awake head-fixed mice. By pairing specific odors with a water reward, we were able to examine how an animal’s associative experience can refine social representations.

Here we report that populations of CA2 neurons initially respond indiscriminately to novel social odors and then with repeated odor exposure rapidly gain the ability to discriminate social odors from two novel mice. Moreover, following training in an operant-conditioning associative reward-learning task, the discrimination of social odors through these representations is enhanced. Finally, whereas populations of CA2 neurons respond to both social and non-social odors, the geometry of these neural representations suggests that there is an abstract code that enables CA2 firing to distinguish between social versus non-social odors. Thus, similar to hippocampal spatially-selective firing (e.g., place cells), distinct CA2 representations of ‘social space’ emerge rapidly and then are refined through subsequent experience.

## Results

### Representations of social odors in CA2

To record specifically from CA2 pyramidal neurons, we injected Cre-dependent adeno-associated virus (AAV) expressing the calcium sensor GCaMP6s (Chen et al., 2013) in dorsal CA2 of the Amigo2-Cre driver line (Hitti and Siegelbaum, 2014) and chronically implanted a cranial window (Dombeck et al., 2010) on top of dorsal CA2 (Figure 1A). Posthoc co-labeling with a marker for CA2 confirmed the specificity of expression (Figure 1B). In the field of view of an in-vivo recording, CA2 pyramidal neurons typically span the diagonal of the imaging plane (Figure 1C) and we identified on average 115 mean ± 21 STD pyramidal neurons per recording session. We head-fixed subject mice under a two-photon microscope and presented the volatile fraction of two different urine samples from age-matched individuals or blank air as control via a custom-built olfactometer while recording activity from CA2 (Figure 1D). A single trial, including either an odor or blank presentation for 1 second, lasted 20 seconds and each stimulus was repeated 20 times, resulting in a total of 60 randomly ordered trials (Figure 1E).

**Figure 1.**
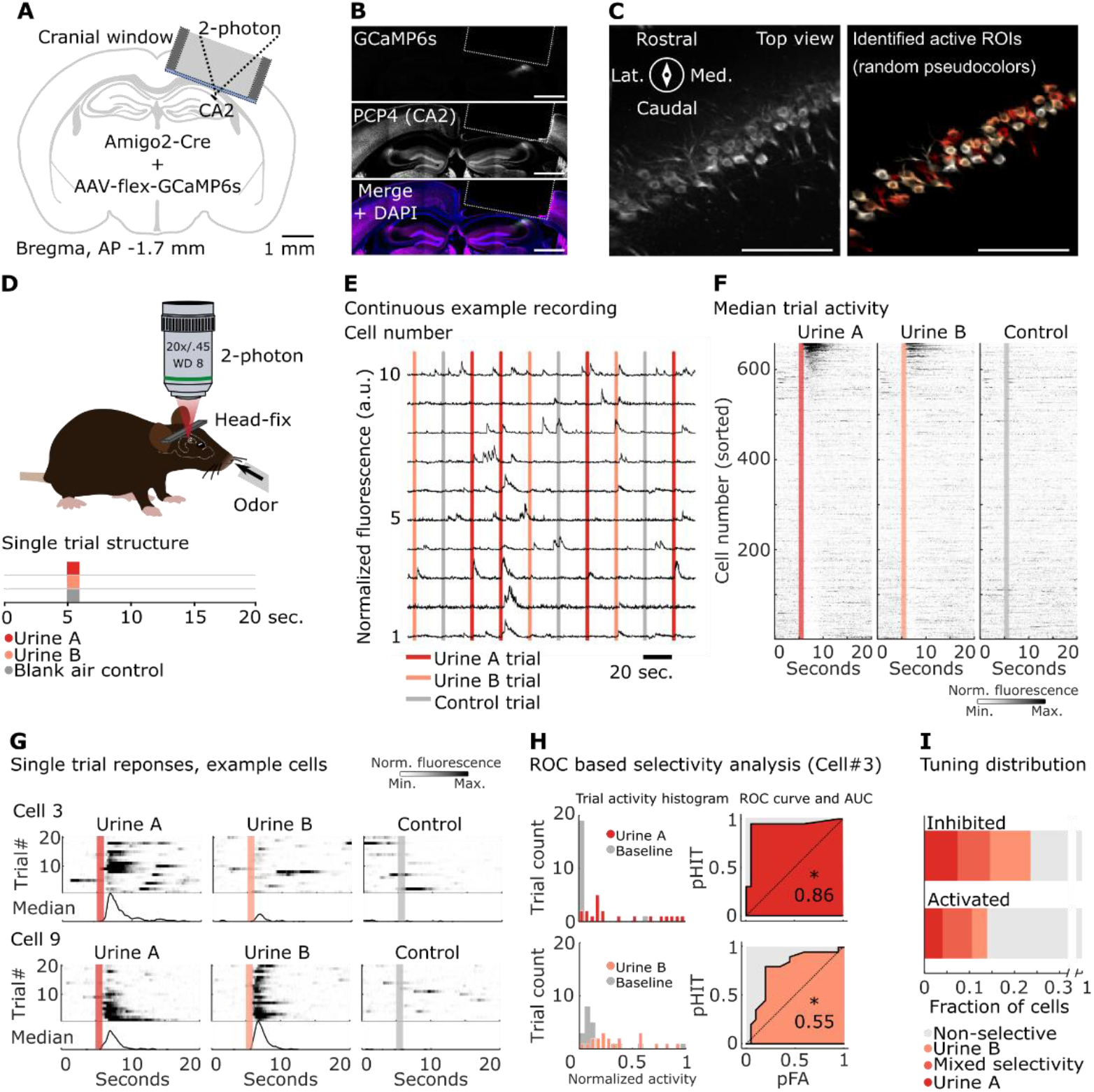
Multiphoton imaging reveals specific tuning to social odors in CA2 PNs. (A) AAV mediated GCaMP6s expression targeted to CA2 PNs using the Amigo2-Cre driver line. An angled cranial window was implanted on top of the alveus and positioned over CA2 for multiphoton imaging in vivo. (B) Confocal image of a brain section from an example animal. Counter-staining with the CA2 marker PCP4 (middle) confirms the specific expression of GCaMP6s (top) in CA2 PNs. Scale bar: 1 mm. (C) Left, in vivo two-photon image (maximum intensity projection) from a representative field of view showing GCaMP6s fluorescence in cell somata and dendrites of CA2 PNs. Right, pseudo-colored cell masks (ROIs) of identified active cells during a single recording session. Lat.: Lateral, Med.: Medial. Scale bar: 100 µm. (D) Experimental design for imaging during passive odor presentation. Head-fixed, immobilized awake mice were randomly presented with one of two different social odors – urine from two unfamiliar mice – or blank air control. A total of 60 trials, 20 trials per odor or control, were presented. ITI: inter-trial interval. (E) Ten simultaneously recorded example cells. Continuous normalized fluorescent signals (black traces) with overlaid timing of random odor/blank air presentations. (F) Median activity of all cells recorded for odor and control presentations (n = 659; 7 animals). Colored/grey bars indicate the timing of stimuli. Cells are sorted by ascending average odor response zero to three seconds after odor onset in Urine A condition. (G) Single trial responses of two example cells, sorted by stimulus condition. (H) Calculation of CA2 PN stimulus selectivity. Left, Single trial activity histograms for cell 3 (from (G)). Top, responses to control (grey bars) or urine A (dark red bars); Bottom, responses to control or urine B (bright red bars). Right, ROC analysis for odor discrimination compared to control for cell 3. Top, Odor A; Bottom, Odor B. Probability of true positive (pHit) plotted versus probability of false alarm (pFA). Shaded regions show the area under the ROC curve (AUC). Significance of selectivity (*, p < 0.05; n.s., non-significant) was determined through sampling under the null-hypothesis (no difference between baseline and post-odor activity). (I) Distribution of odor response selectivity across cells from ROC analysis. Top bars, Cells with significantly lower activity in response to urine A, urine B, or to either A or B (mixed selectivity) relative to control. Bottom bars, Cells with significantly greater activity in response to urine A, urine B, or to either A or B (mixed selectivity) relative to control.

Sorting trials by their respective stimulus type and plotting the median trial activity of individual cells as a function of a single trial time course revealed subsets of cells with increased or inhibited activity, relative to baseline, to either one or both of the two urine samples. In contrast, control air-only trials did not elicit a response (Figure 1F). The single-trial analysis further confirmed robust odor-evoked responses in a fraction of cells. Some cells showed consistent selective activation in response to one odor over another, despite a dynamic response pattern for individual odors across a single session (Figure 1G, cell 3). Other cells showed a more complex selectivity pattern that changed tuning to one or another odor over the course of a single recording session (Figure 1G, cell 9).

For a more precise measure of tuning properties, we employed receiver operating characteristic (ROC) analysis (Green and Swets, 1966; Najafi et al., 2020). For each neuron, we compared the trial activity distribution before and after stimulus presentation (Figure 1H) and a neuron was identified as odor ‘activated’ when the area under the ROC curve (AUC) differed significantly (p < 0.05) from a shuffled distribution. We found 13.9% of pyramidal neurons were activated by one or more social odors (3.9% were activated by urine A only, 3.3% by urine B only, and 6.7% by both A and B). A slightly larger fraction of cells, 24.2%, were significantly inhibited in response to odor (with 7.9% inhibited by odor A only, 9.6% by odor B only, and 6.7% of cells inhibited by both A and B) (Figure 1I, Figure S1B).

### CA2 social stimulus representations change with their degree of novelty

The structured response of pyramidal neurons in CA2 to social odor presentation of single cells indicates that CA2 may encode the identity of a given social olfactory stimulus. To test this hypothesis, we examined the CA2 pyramidal neuron population activity. We first constructed pseudo-population vectors that contained the individual trial activity averaged over three seconds after odor presentation for each cell and performed principal component analysis (PCA) on the cell-by-cell activity covariance matrix (Figure 2A). The first three principal components captured 74.7% of the variance. Visualization of the trial activity revealed that the responses to the two odors and blank control formed distinct clusters in the low-dimensional PCA space. Control versus social odor responses were distinguished by the magnitude of their first principal component, whereas the responses to the two urine samples were distinguished by the magnitudes of their second and third components (third axis not shown). These representations were dynamic as we observed differences in the population responses to the same odor presented in an early compared to late trial within a given twenty-minute session, with responses becoming more distinct as the trials progressed.

**Figure 2.**
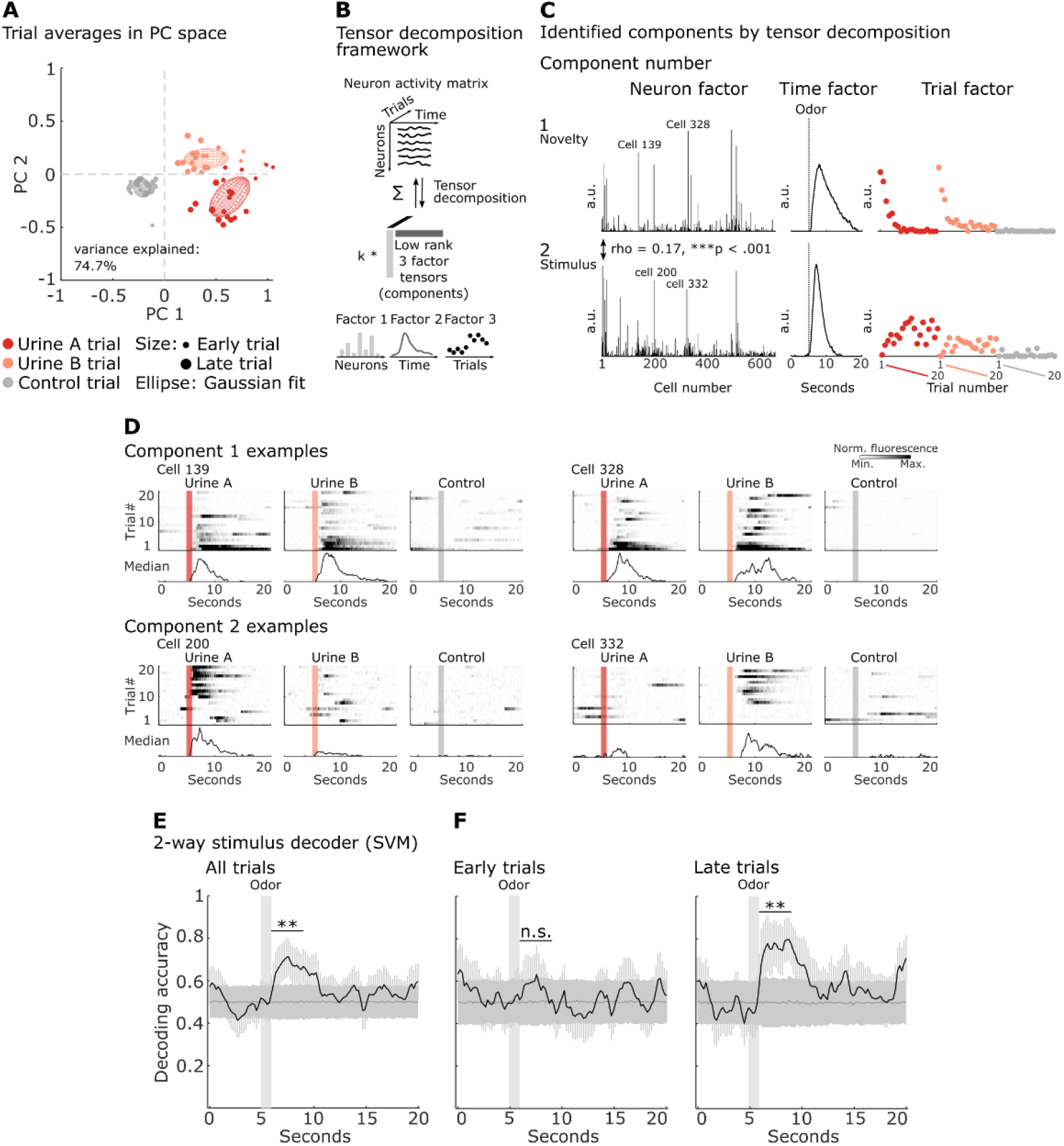
CA2 decodes social odors with high accuracy. (A) First two PCA components of CA2 population trial responses in a three-second window after stimulus presentation to social odors or blank air control. Individual dots represent PCA projections of single trials. A two-dimensional Gaussian model is fit to each stimulus trial data (ellipses; mean ± 1 SD). The size of the dots indicates the time of a trial within a session: small dots, early trials; large dots, late trials. Units: normalized fluorescence. (B) Schematic of the tensor decomposition approach. Neuronal activity across different cells and trials is organized in a three-dimensional data matrix. Tensor decomposition approximates the data as the sum of k low-dimensional components that are the outer product of three vectors (factors). These factors can be interpreted biologically as a neuronal ensemble (factor 1), with a particular response time course during a trial (factor 2) that follows a specific trajectory across trials (factor 3). (C) The original data were decomposed into two reproducible low-dimensional components, each containing a neuron ensemble (left column), temporal factor (middle column), and trial factor (right column), based on a 12-component model. (D) Single trial raster plot responses and median response of two example cells with high neuron factor weights in each component (identified in C by their number). The random succession of odor trials is re-arranged for clarity for each stimulus condition, from early to late trials within each stimulus block. Color code as in (A). (E) Left, binary stimulus classification accuracy (black trace) or chance accuracy (dark grey trace) for decoding of social odor identity (urine A versus urine B) as a function of time during odor trials. Results are the average over 1,000 repetitions of random sub-samples of 200 cells; **, p < 0.01; empirical p-value based on the distribution of paired differences between the subsampled signal and chance distributions. (F) Average binary stimulus classification accuracy for early and late trials. Decoding was run separately using either the first (left) or second half (right) of the total odor trials. 1,000 random sub-samples, sub-sample size: 200. Color code as in (E). SVM: support vector machine decoder; **, p < 0.01, n.s., non-significant. Panels (A, C, E, F) are based on pseudo-population data, constructed from 659 cells from seven animals; five-fold cross-validation was used for all decoding; error bars represent SEM.

As PCA has limited capabilities to capture both, short- and long-term temporal dynamics (within and across trials, respectively), we used tensor decomposition (TCA) of a matrix factored in three dimensions: 1. The individual neurons of the population; 2. Time during a single trial; 3. The individual trials during a session (Williams et al., 2018). A TCA fit of a model of low-dimensional latent components to the matrix yielded two prominent components composed of distinct but overlapping subsets of neurons (in contrast to PCA, TCA does not enforce orthogonality between components; Spearman rank correlation of neuron factors, rho = 0.17, p < 0.001).

The first component showed a time-factor weight increase time-locked to odor onset for both odors. Moreover, the weight of this component was greatest in the early trials of a session, and then rapidly decreased for both odors. This suggests that this component represents the novelty of the stimulus; control blank trials failed to activate this component (Figure 2C, top).

The second component comprised a different, yet slightly overlapping set of cells that appeared to reflect an emerging differential response to the two odors. Although the weights of the second component displayed a similar within-trial time course compared to the ‘novelty component’, the trial factor weights of the second component slowly increased during a session. Moreover, whereas both social odors elicited similar trial factor weights in the ‘novelty component’, the pattern of neuronal activation between odors differed from one another in the second component, with slightly lower activation weights for odor B. This suggests that the second component reflects an emerging discrimination in the population response to the two social odors during the trials of a single session (Figure 2C, bottom).

The TCA decomposition into novelty and discrimination classes of neuronal responses was evident in the responses of single neurons that had high neuron weight factors for a particular component. Thus, neurons in the novelty component exhibited non-selective but strong responses to both odors in early but not late trials (Figure 2D, top), whereas neurons with high weights in the discrimination stimulus component showed responses whose strength and selectivity for a given social odor increased over the course of a session (Figure 2D, bottom). In summary, both PCA and TCA models indicate that overlapping subsets of CA2 neurons can distinguish social odor identity *and* detect the degree of novelty of a stimulus.

To further explore the information content of the CA2 pyramidal neuron population in a manner that did not require dimensionality reduction, we turned to neural decoding analysis. To address the question of stimulus identity, we trained and tested a binary linear classifier (support vector machine (SVM); (Hofmann et al., 2008)) on single-trial responses to both urine samples as a function of single-trial time, using random sub-samples of 200 cells from our pseudo-population (Figure 2E). Before odor onset, the classifier performed at chance. Once the odor reached the nose port, decoding accuracy sharply rose significantly above chance and then decayed back to chance during the inter-trial interval (pairwise difference signal minus chance accuracy = 0.16 mean ± 0.06 SEM; p_empirical_ = 0.006; 1000 sub-samples).

We obtained complementary results using a second classifier based on the cosine distance of the population vectors. In comparison to the regular implementation of a binary SVM, this classifier is able to separate all three stimulus conditions (control plus the two odor trials). This confirmed the significant stimulus decoding accuracy (pairwise difference signal minus chance accuracy: 0.35 mean ± 0.05 SEM; p_empirical_ = < 0.001, 1000 sub-samples). Further, the classifier’s high true positive rate for control trials supported CA2’s ability to discriminate odor from blank air control trials, which was suggested earlier by the PCA model (control trials: true positive rate = 0.79, chance level = 0.33; Figure S2A).

As the PCA and TCA analyses indicated that experience during a session increased odor discrimination, we examined how decoding accuracy changed during the session. Indeed we found that in early trials the stimulus decoding failed to rise above chance level (pairwise difference signal minus chance accuracy = 0.06 mean ± 0.08 SEM; p_empirical_ = 0.2; 1000 sub- samples; Figure 2F, left; Figure S2B), whereas in late trials the average decoding accuracy after odor presentation significantly exceeded chance levels, and surpassed the decoding accuracy using all trials in a session (pairwise difference signal minus chance accuracy = 0.24 mean ± 0.09 SEM; p_empirical_ = 0.004; 1000 sub-samples; Figure 2F, right; Figure S2B). The change in neural activity across a session allowed a binary decoder to successfully classify trials from either the first or the second half of the session irrespective of stimulus identity (pairwise difference signal minus chance accuracy = 0.13 mean ± 0.04 SEM; p_empirical_ = 0.002; 1000 sub- samples; Figure S2C, left panel). As a temporal control, we trained a separate decoder to classify alternating trials, which did not perform better than chance (pairwise difference signal minus chance accuracy = -0.03 mean ± 0.03 SEM; p_empirical_ = 0.8; 1000 sub-samples; Figure S2C, middle panel).

### CA2 social odor population code is driven by a subset of odor-selective neurons

The significant decoding of odor identity in CA2 can be explained either by the highly selective responses of a subpopulation of cells or a population-based response of cells with weak, mixed selectivity (Stefanini et al., 2020). To distinguish among these possibilities, we performed a decoding analysis using random subsamples of 200 neurons (out of the total population of 659) that excluded the 65 neurons found by ROC analysis to be activated by the social odor from one mouse to a significantly larger extent compared to the social odor from the other mouse. We focused our analysis on the trials during the last half of the session when decoding accuracy was greatest. Decoding with only the non-selective cells failed to rise above chance level (pairwise difference signal minus chance accuracy = 0.05 mean ± 0.06 SEM; p_empirical_ = 0.2; 1000 sub-samples; Figure S2B). On the other hand, subsamples from the pool of selective cells showed saturation of the decoding accuracy with only 20 cells (titration data not shown), a tenth of the initial subsample size of 200, and led to slightly better decoding accuracy compared with results from the whole population (pairwise difference signal minus chance accuracy = 0.28 mean ± 0.07 SEM; p_empirical_ = 0.001; 1000 sub-samples; Figure S2B). Thus decoding accuracy appears largely driven by a subset of odor-selective cells that gain their selectivity during experience over the multiple trials of a single session.

### Associative learning during a social odor based reward-pairing Go/No-Go task

Having observed the acquisition of distinct representations of novel social odors during experience, we next examined how these representations might be affected by association of one odor with positive valence. Following the initial odor presentation session described above, we presented the same two social odors in a trial-based Go/No-Go associative reward learning task (Li et al., 2017). A single trial lasted 20 seconds and consisted of a baseline, followed by a one-second-long odor or blank presentation, a 3-s delay for odor sampling, a 1-s LED cue window during which the mouse was required to lick at a spout to obtain a water reward, followed by a 10-s inter-trial interval. Mice were trained to associate odor A with reward, with urine B and blank air presentations unrewarded (Figure 3A). Mice rapidly learned to lick following odor A and to withhold licking to either odor B or blank air, reaching criteria of 80% correct trials after 2-5 days of training (Figure 3B, C). We imaged CA2 activity to odor presentation before training (as described in Figures 1, 2 above), during training, and for two days after reaching criterion.

**Figure 3.**
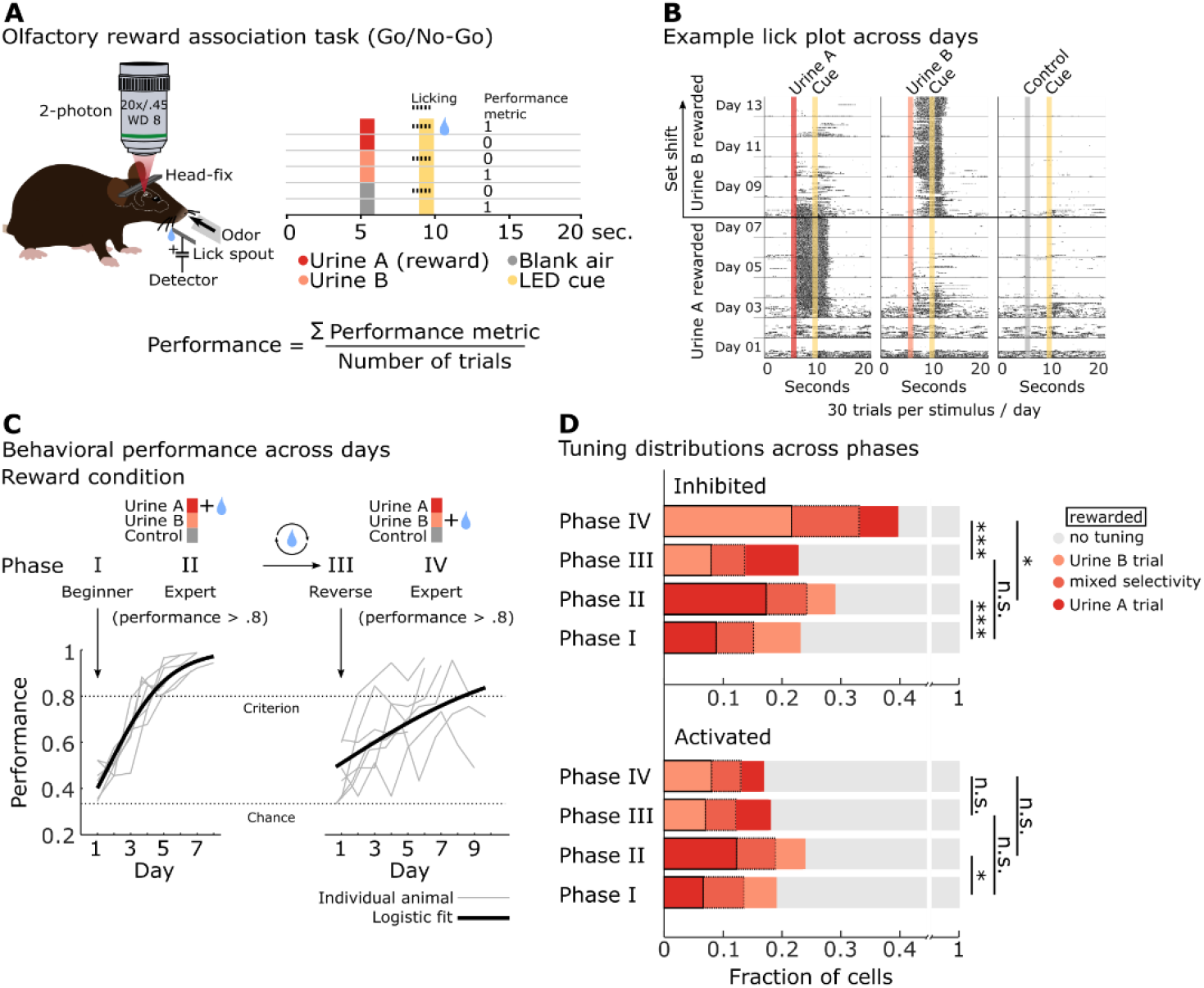
Mice learn flexible social odor-reward associations in a Go/No-Go odor-reward association task. (A) Diagram of behavioral setup, the task structure, and the outcome metric to calculate a performance index. See text for explanations. (B) Example lick plot from an individual animal performing the task across 13 days. Each day, 90 randomly ordered urine and control (blank air) trials were presented to the animal (30 trials per condition). In the plot, random single-day trials are split across stimulus columns and re-sorted from early trials at the bottom to late trials at the top of a single day. Days are then stacked on top of one another. Rows represent single trials; black tick marks are individual lick contacts with the spout; colored / grey bars indicate stimulus timing, yellow bars show the cue window; horizontal grey lines separate individual days, the thick black horizontal line indicates the time of set-shift in reward contingency. (C) CA2 PN activity was recorded over the course of learning and different sessions were classified into four different phases based on the mouse’s behavioral performance: Phase I, the first day of training prior to learning the task; Phase II, two more consecutive days after a mouse reached criterion (> 80% correct trials); Phase III, the first day of reward reversal, prior to full learning : the formerly non-rewarded urine B was now associated with reward and odor A was unrewarded; Phase IV, comprises a single session after an animal learned the reversed contingencies (>80% correct trials). Lower panels show progressive learning of the task by individual animals (grey lines), either before (left lower panel) or after (right lower panel) reward reversal. The increase in performance as a function of trials is fitted by a logistic function (black line). Performance levels: criterion 80%, chance 33%. (D) ROC based tuning distribution of significantly activated or inhibited CA2 PNs (relative to activity during pre-stimulus baseline) across the different phases of the experiment. Significant selectivity with p < 0.05 was determined through sampling under the null-hypothesis. Bonferroni-corrected p-values for the Fisher exact test: *, p < 0.006; ***, p < 0.0001, n.s., non-significant.

Besides being associative, social behavior is also adaptive and often requires flexible adjustments of representation. We thus switched reward contingency after mice reached stable performance, rewarding odor B instead of A, and followed re-learning of the task under the new reward contingency (Figure 3B, day 8-day 13; Figure 3C, top, Phase III & IV). All mice (n = 7) learned to associate odor A with reward within five days and a logistic fit to model their group-wise learning crossed the criterion level at day four (Figure 3C, bottom left). Reversal learning proved more challenging; only four out of seven mice learned the contingency switch above criterion, resulting in a rightward shift of the logistic model crossing the criterion level (day eight).

### CA2 social odor tuning under reward learning

To explore how reward learning may alter social odor representations, we pooled cells from all recorded animals during each of the four phases of the experiment (see Figure 3C for details) and performed ROC analysis to determine odor response selectivity of single cells (Figure 3D). Within the population of cells that responded to either odor A or B, the fraction that responded to the rewarded odor increased after both initial (from 0.35 to 0.51) and reversal learning (from 0.39 to 0.48), at the expense of a reduction in the fraction responding to the non-rewarded odor, during initial (0.29 to 0.21) and reversal (0.33 to 0.23) learning. The fraction of cells activated by both odors stayed constant, around 0.30. Interestingly the most dynamic changes were observed in the fraction of cells inhibited by rewarded odor, relative to the population inhibited by either rewarded or unrewarded odors Here, the fraction selectively inhibited by rewarded odor increased during initial learning from 0.38 to 0.60 and from 0.35 to 0.55 during reversal learning. The fraction selective to the non-rewarded odor decreased from 0.34 to 0.16 during initial learning and from 0.40 to 0.16 during reversal learning. The mixed fraction remained unchanged, around 0.26. A Fisher exact test showed a significant effect between experimental phase and tuning distribution for the transition from Phase I to Phase II in activated cells; in the inhibited fractions this effect was significant for both the transition between Phase I and Phase II, Phase III and Phase IV and also between Phase II and Phase III.

### Tuning of CA2 pyramidal cells reflects stimulus identity, not behavioral choice

One caveat in our task design is that odor presentation is always connected with a choice: to lick or not to lick. Any tuning we observed on the single-cell level might therefore represent either stimulus tuning, or the decision that the animal is about to make, or both. To clarify the nature of the tuning, we leveraged behavioral error trials in the early learning phase (Feierstein et al., 2006) by comparing tuning selectivity when the animal made a correct choice, e.g., licking after odor A presentation, with the tuning properties in error trials, e.g. failure to lick to odor A. We observed a highly significant correlation in selectivity index to a given odor in a plot of error versus correct trials (Figure S3A; Spearman rank correlation ρ = 0.70, p < 0.001; 44 from 677 cells total are significantly tuned in both correct and error trials; n = 6 animals).

Together with the observed tuning during non-rewarded odor presentation, where no choice component existed, we conclude that CA2 neuron tuning is stimulus-dependent and not significantly modulated by the animal’s behavioral choice.

### CA2 forms flexible social odor-reward associations, which improve decoding of stimulus identity

To examine how the dynamic changes at the single-cell level influence the CA2 population code, we first performed PCA analysis of the pseudo-population, compiled for each of the four phases separately (Figure 4A). The PCA plot on the first day of learning resembled that of the non-rewarded presentation: distinct clusters for odors A and B were separated from mock trials along the principal axis one, whereas the principal axis two separated the odor clusters. In line with the odors being already familiar at this point, no novelty dependence (trial number) was apparent (average behavior performance: 0.43 ± 0.03 SEM; n = 6). Learning the reward association (> 0.8 performance) induced striking changes in the cluster distribution, with the rewarded odor becoming well separated from both control (blank air) and the unrewarded odor along the first principal axis. Principal axis two still separated all three trial types.

**Figure 4.**
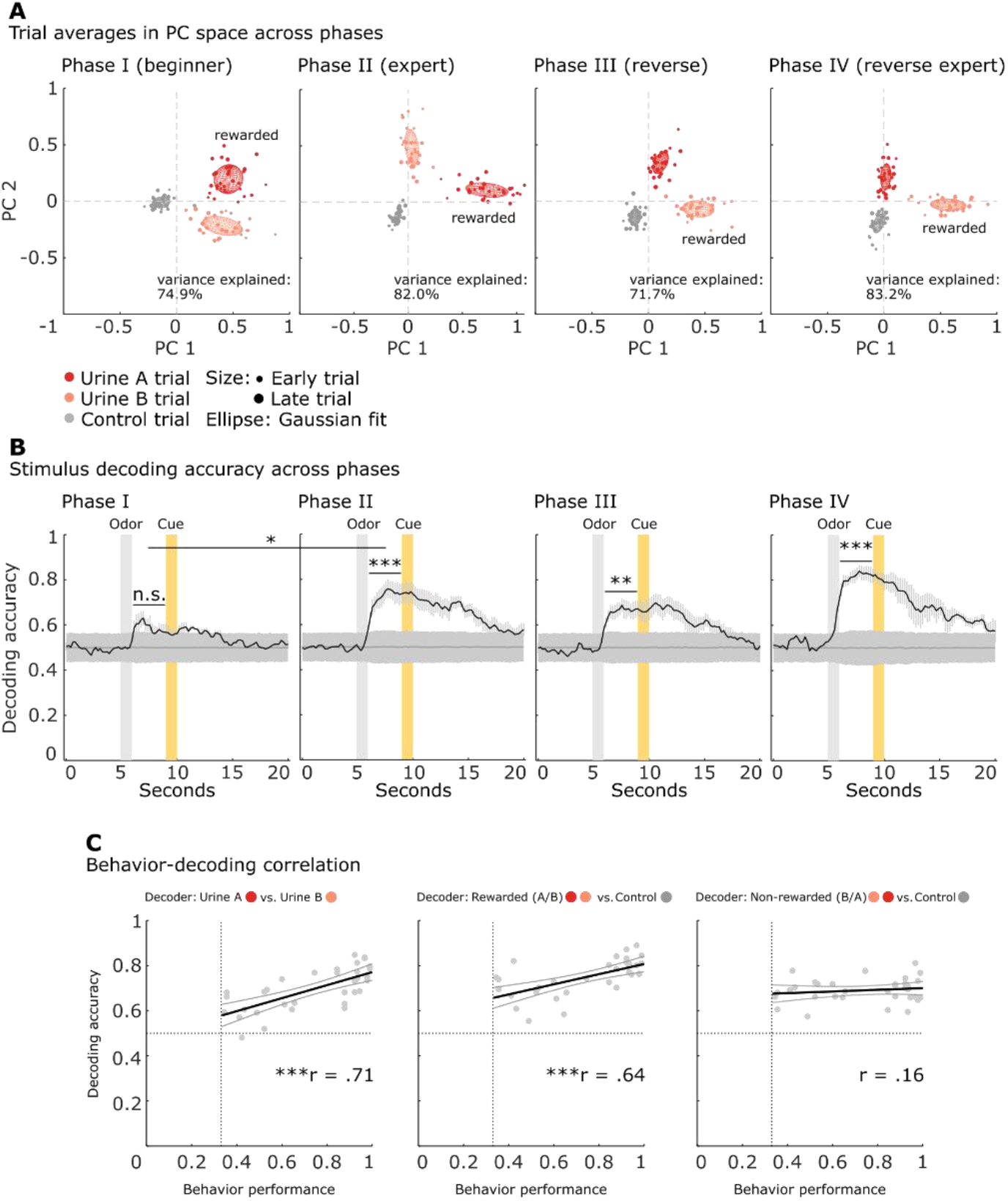
Odor discrimination is enhanced following training in the Go/No-Go task. (A) PCA representation of CA2 population trial responses in a three-second window after stimulus presentation to social odors or blank air control across the different phases of the odor-reward association task. Units: normalized fluorescence. (B) Decoding accuracy across the different phases of the task. Linear decoders were trained and tested, separately for each session and mouse across the different phases. Multiple random sub-samples (70 cells, 1,000 samples per mouse) of simultaneously recorded cells were chosen and a decoder was trained and tested to classify the two different urine samples. Significance within session was determined in the three second delay window after odor presentation by calculating an empirical p-value using the pairwise difference distribution between signal and chance distributions; significance of the increase of decoding accuracy between Phase I and Phase II was determined using repeated-measure ANOVA; n.s., non-significant; *, p < 0.05, **, p < 0.01; ***, p < 0.001. (C) Behavior-decoding correlation. The session based performance index of individual animals was correlated with their respective decoding accuracy in the same session. Decoders were either trained to distinguish between the two urine samples (left), the rewarded odors (urine A before the set shift and urine B thereafter) and the control trials (middle), or the non-rewarded odors and the control trials (left). Individual sessions of six animals across the four phases are plotted as grey data points. A linear regression line (black thick line) and its 95% CI (thin lines) are overlaid; ***, p < 0.001.

Remarkably, on the first day of reward reversal when learning was not yet complete (average performance: 0.50 ± 0.04 SEM; n = 6), the rewarded and unrewarded odor clusters switched positions in PCA space, which then became further refined in those animals (n = 3) that reached criterion when reversal learning was complete.

Both dynamic changes on the single-cell level as well as the PCA analysis suggest that CA2 flexibly incorporates valence in existing stimulus representations. We hypothesized that this would enhance stimulus classification. To test this, we performed decoding analysis in the full vector space across the different phases. In contrast to the passive odor case, where no learning was involved, mice learned the reward-association task at different rates. To account for inter-individual differences, we trained and tested binary linear decoders for each mouse individually, using subsamples from the simultaneously recorded population and averaged the decoder accuracy across animals. We chose a conservative sub-sample size of 70 cells. This size was slightly less than the lower bound of the number of cells identified across sessions and animals (range: 87-168) to enable us to detect subtle differences in decoding performance among the different phases, which might be obscured by saturating performance levels with larger number of cells.

Using this smaller subsample (compared to the 200 cells used for passive odor decoding), the linear classifier failed to decode odor identity above chance levels during the initial passive odor presentation (measured minus chance decoding accuracy = 0.09 ± 0.08; mean ± SEM; p_empirical_ = 0.2; Figure 4B, Phase I; Figure S5B). In contrast, decoding accuracy became highly significant after associative learning (measured minus chance accuracy = 0.21 ± 0.09; p_empirical_ = <0.001; Figure 4B, Phase II; Figure S5B). The lack of significant stimulus decoding is a reflection of the smaller subsample size, as stimulus decoding using 200 cells from a pseudo-population composed of all cells from all animals in Phase I, reached significance (measured minus chance decoding accuracy = 0.13 ± 0.05; mean ± SEM; p_empirical_ < 0.001; data not shown).

To explicitly test for learning induced changes in CA2, we performed a repeated-measure ANOVA focusing on Phase I and the first day of Phase II as not all animals learned the ensuing reversed reward condition. The test showed a significant effect of learning on decoding accuracy (F(1, 5) = 7.8, p = 0.04). On the first day of training following the switch in reward contingency, there was a drop in decoder performance, however, accuracy was still better than chance (measured minus chance accuracy = 0.12 ± 0.07; p_empirical_ = 0.001; Figure 4B, Phase III; Figure S5B). In those animals that reached criterion after reward reversal, decoding accuracy attained its high level seen during the initial reward-association training (measured minus chance accuracy = 0.31 mean ± 0.06 SEM; p_empirical_ < 0.001; Figure 4B, Phase IV). These decoder results suggest that learning improves decoding accuracy.

To test for a dependency between behavioral performance and decoding accuracy, we plotted decoding accuracy versus behavioral performance for all animals during all sessions and calculated the Pearson correlation coefficient, including animals that did not reach criterion after the contingency switch. We observed a highly significant linear relation between the accuracy with which a linear classifier decoded the two odors and behavioral performance (r = 0.71, p<0.001; Figure 4C, left panel). We also saw a significant relationship between decoding of the rewarded odor (either A or B) versus blank air control and behavior (r = 0.64, p<0.001; Figure 4C, middle panel). However, there was no significant relationship between decoding of non-rewarded odor (either A or B) and the blank air control (r = 0.16, p = 0.4; Figure 4C, right panel), which indicates that the learning dynamic reflects the increased incorporation of reward information into the social odor representation, which might reflect secondary effects other than reward itself, such as increased attention towards the rewarded stimulus.

### Representational drift in CA2

Two-photon microscopy allowed us to reliably track the same cells over different days (Figure S3B). We compared tuning properties of individual cells across sessions (Chéreau et al., 2020) by calculating a selectivity index for each cell using ROC analysis in the different sessions (Figure S4A). For each cell, we compared pairs of sessions by plotting its selectivity index measured on a given day versus its selectivity index on a preceding or subsequent session. Cells that were significantly tuned in at least one session are shown as black dots (Figure S4B-D). We classified cells into four groups: cells that lost their selectivity from one day to the next; cells that gained selectivity; cells that retained their selectivity between sessions; cells that reversed their selectivity. We found a notable dynamic among cells, both during learning (Figure S4B) and once the animal reached criterion (Figure S4C and D), with a majority of cells either gaining or losing selectivity across sessions. After learning, however, the fraction of cells that retained their selectivity increased, which resulted in a significant positive correlation in the total fraction of significant cells across the expert phase (Figure S4C and D).

We also compared the conditional probability of a class to its theoretically expected probability in the case of independent sessions and found significant results for cells losing their tuning in all three cases, which indicated a predisposition of once active cells to lose their tuning in a later session.

The high degree of fluidity motivated us to test how decoding across days might be affected. We focused on the expert phase, when animals showed stable behavioral performance above criterion and trained a decoder to distinguish urine A from urine B and then tested the decoder on the activity of the same cells in a subsequent session (Figure S5A and B). The decoding with random subsamples of 70 cells showed significant decoding between successive days whereas decoding across a two-day interval dropped to chance levels. The conservative subsample size and the linear decoding regime most likely result in a lower bound estimate of decoding accuracy across days. Nevertheless, these results paint a dynamic picture of CA2’s tuning properties despite a consistently high information content within a given session (Figure S5B).

### Inactivation of CA2 impairs olfactory associative learning using social odors

The significant correlation between the CA2 decoding accuracy and the degree of performance in the social-odor-reward association task led us to test for a causal role of CA2 in forming the odor-reward association. We bilaterally injected a separate cohort of Amigo2-Cre mice with AAVs that express in a Cre-dependent manner either the inhibitory opsin archaerhodopsin 3.0 (eArch3.0) or the fluorescent marker mCherry, as a control, and implanted optical fibers on top of the alveus right above dorsal CA2 to silence pyramidal neuron somata (Figure 5A). We included only animals in the analysis where we confirmed posthoc a confined viral expression and correct fiber placement (Figure 5B). We shined green light (532 nm) in each trial for eight out of twenty seconds, starting one second before odor sampling and stopping two seconds after the cue window had ended (Figure 5C). Control animals expressing mCherry steadily increased their performance and crossed on average the criterion performance of 80% correct association trials by day four (Figure 5D, left; Figure 5E). However, in mice expressing eArch3.0 in CA2, optogenetic silencing significantly impaired the learning process; on average, the mice never reached the 80% criterion and the fit of a three-parameter logistic function to model the learning progression revealed a significant decrease in maximal performance in eArch3.0 mice compared to controls (control: coefficientmax = 0.910, n = 5 mice; eArch3.0: coefficientmax = 0.743, n = 4 mice; p_empirical_ = 0.007, 5000 permutations; Figure 5E).

**Figure 5.**
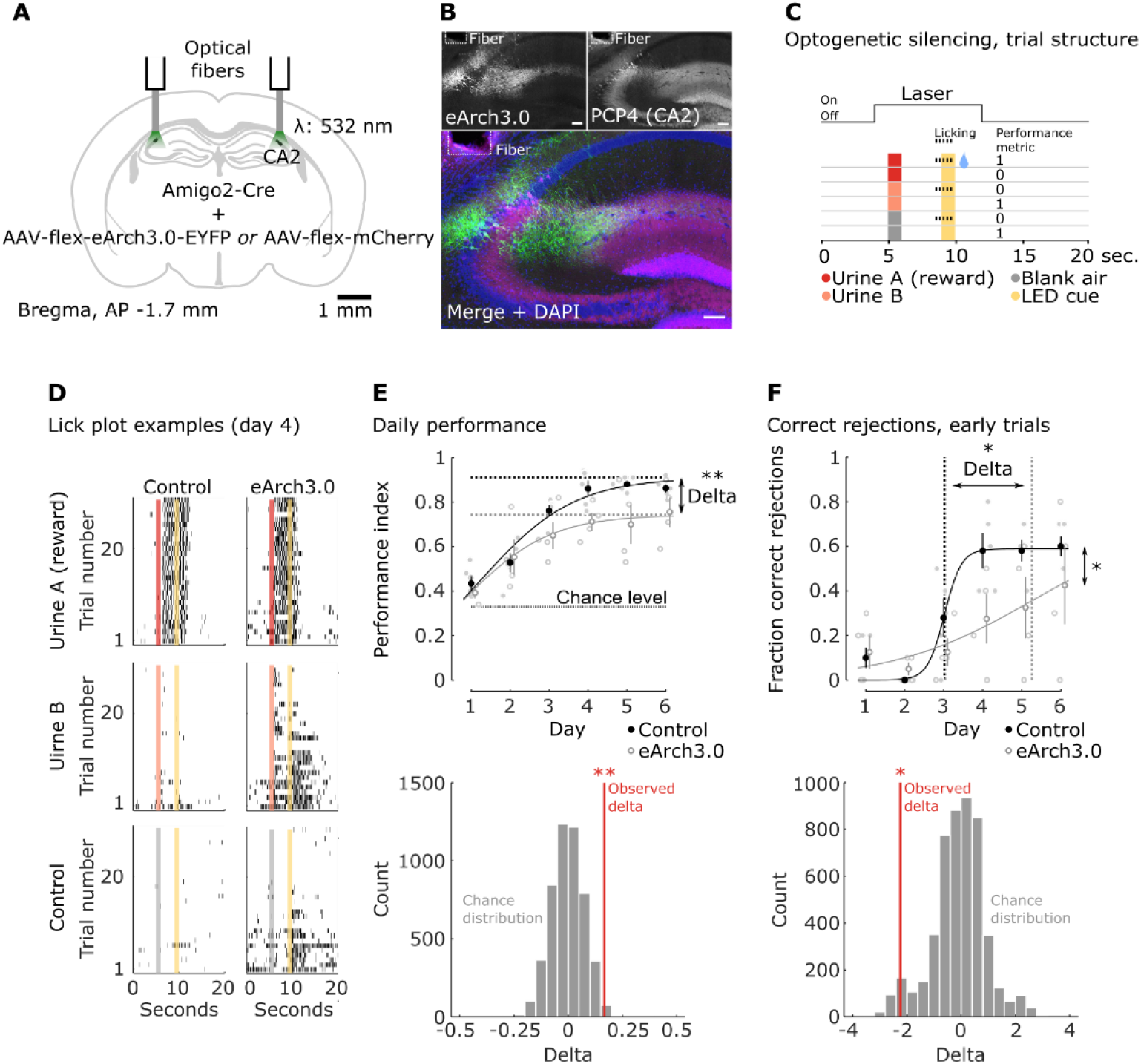
Optogenetic silencing of CA2 impairs the learning of social odor-reward associations. (A) AAV mediated expression of either Archaerhodopsin (eArch3) or mCherry control targeted to CA2 PNs using the Amigo2-Cre driver line. Optical fibers positioned on top of CA2 allow acute silencing of CA2 PN activity during behavior. (B) Confocal image of a brain slice from an example animal in the eArch3-group. Counter- staining with a marker for CA2 (PCP4, top right panel) confirms concise expression of eArch3 in CA2. (C) Schematic of the experimental paradigm. In a given trial, laser light was provided starting one second before odor onset until two seconds after the cue window/reward presentation (total of 8 s per 20 s trial). Mice of both groups performed 90 trials per session, 30 trials per stimulus. The course of learning was observed over six sessions, with one session per day. (D) Example lick plots on day four of the task from control and eArch3 expressing animals (individual columns). Randomly presented odors are re-sorted and plotted as a block within a column. (E) Upper panel, the performance of individual animals (light circles: filled = controls, open = eArch3) plotted as a function of training day. Mean ± SEM (larger circles); a three parameter logistic regression model is fit to the data of each group. Delta represents the difference of the plateau parameter of fitted models to each group respectively. Significance is determined using a permutation test (lower panel); **, p < 0.01. (F) Fraction of correct rejections in the first 30 trials of a given session. Impaired learning is reflected by a significant shift of the midpoint/inflection point of a logistic model on top of an overall effect of group on the fraction of correct rejections (repeated-measure ANOVA, *, p < 0.05); the difference in delta is determined using a permutation test (lower panel); *, p < 0.05.

Closer inspection of the licking behavior in the silenced group revealed that CA2 silencing caused the most striking behavioral impairment during the early trials of a given session, where mice need to recall memory of the task contingency from the previous day’s session (Figure 5D, right). This is consistent with the role of hippocampus in mediating long-term association memory (Eichenbaum, 2017). Furthermore, the effect of CA2 silencing was particularly pronounced when we focused on the animal’s learning to withhold licking to the unrewarded odor. This is likely because the default behavior of the animal prior to learning was to lick to the light cue independent of any odor so that learning the task requires the learned suppression of licking to the unrewarded odor. We observed that CA2 silencing produced a marked rightward shift of the inflection point of a logistic fit to a plot of performance to withhold licking to odor B (Park et al., 2021) (control: coefficientinflection = 3.02 days, n = 5 mice; eArch3.0: coefficientinflection = 5.26 days, n = 4 mice; p_empirical_ = 0.04, 5000 permutations; Figure 5F). In addition, a repeated-measure ANOVA indicated a significant effect of the experimental group variable on performance with no significant interaction (Fgroup(1, 7) = 6.3, p = 0.04; Fgroup*day(5, 35) = 1.6, p = 0.2; Figure 5F). Taken together, these results support a crucial role for CA2 network activity in the acquisition of social olfactory associations.

### CA2 activity decodes social odor identity irrespective of reward association

The enhanced decoding of social odors following reward learning could reflect an increased discriminability of the sensory qualities of the odors themselves, enhanced discriminability based on reward association, or a combination of the two. To explore these possibilities, we trained a group of animals (n = 3) on a set of four novel social odors (urine C-F; Figure 6), two of which were rewarded (urine C and E) and the other two were not (urine D and F). One animal crossed criterion within the first session whereas the other two required two and three sessions. After all animals had crossed criterion (Figure 6B), we recorded CA2 neuronal responses to the four odors. Remarkably, all four odors formed distinct clusters in principal component space (Figure 6C); as in the two-odor case, principle axis one separated rewarded from non-rewarded odors. We also calculated the average cluster distance based on the cosine similarity (the angle between two clusters from the origin) using the first 12 principal components (>90% variance). The overall within-cluster distance to a given odor was small (diagonal values) compared to the across-cluster distances. Non-rewarded odors clustered closer together than rewarded odors, again suggesting, that reward association, or reward expectation, might enhance discrimination.

**Figure 6.**
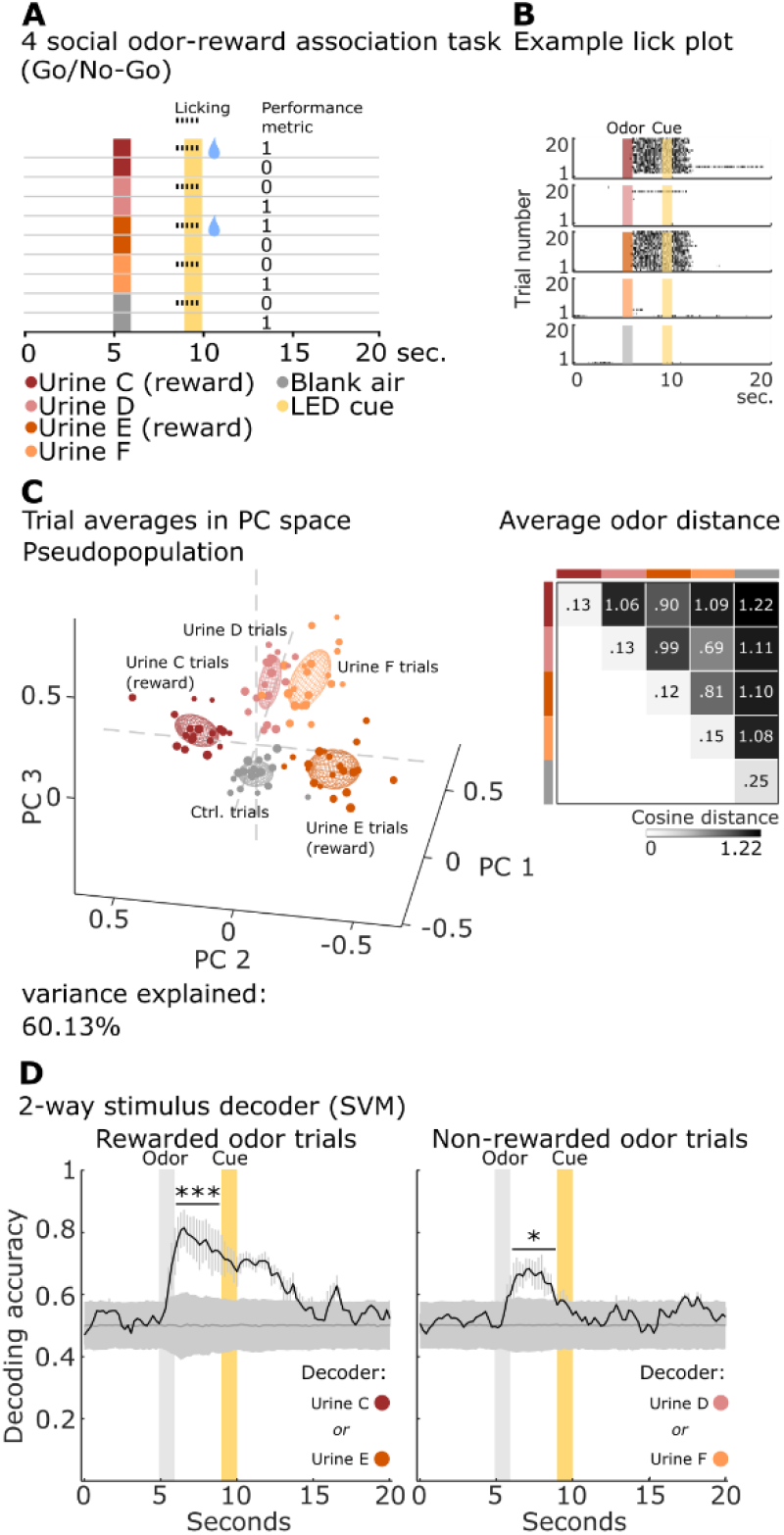
CA2 activity decodes social odor identity irrespective of reward association. (A) Schematic of the four social odor-reward association task and metric to calculate performance. A novel set of four social odors (urine from individual unfamiliar mice) and a blank control are randomly presented during multiphoton imaging. Mice received a water reward when they licked during the cue window after urine C or E were presented. (B) Example lick behavior of a well-trained animal performing above criterion (>80% correct responses). Random trials are re-sorted and plotted in their respective stimulus blocks. Each row represents a single trial; black dash marks indicate licking. Color code as in (A). (C) Left, Three largest PCA components of CA2 pseudo-population trial responses (340 cells from three animals). The activity was averaged across a three-second window after stimulus presentation (social odors or blank air control). Symbols as in Fig 2A, except data fit by three-dimensional ellipse. Right, odor distance in principle component space. Average cosine distance within (diagonal) and across (off-diagonal) stimulus-clusters was calculated using the first 12 principal components, which cover a minimum of 90% variance in the data. (D) Average binary stimulus classification accuracy, excluding blank trials, as a function of time. Decoding of stimuli was run separately within reward category (rewarded odors, left; non-rewarded odors, right). 1,000 random sub-samples with a sub-sample size of 70 cells per animal. Black trace is the average stimulus decoder across three animals; the dark grey trace is chance decoding as a function of time. Error represents SEM. Significance was determined by calculating an empirical p-value based on the paired difference distribution between signal and chance in the three second delay window after odor presentation. Left: decoding among rewarded odors; right: decoding among non- rewarded odors; *, p < 0.05; ***, p < 0.001.

We then trained a binary linear classifier to distinguish odors with their respective reward category (Figure 6D). Both the rewarded odor classifier as well as the non-rewarded one performed significantly better than chance after odor onset. Decoding accuracy in the non-rewarded case ranged somewhat lower than in the rewarded case, however, this difference did not reach statistical significance in the time window after odor onset. In contrast, when we compared the signals in the three seconds following reward delivery, the two decoders differed significantly, with reward decoding outperforming non-reward decoding, underlining the finding that CA2 has access to reward information (Figure S3C). This complimentary and independent set of results unequivocally shows CA2’s capacity to classify social odor identity of four distinct individuals in a manner that is modulated and fine-tuned by reward association, a prerequisite of social discrimination and social value associations.

### CA2 activity is modulated by both social and non-social odors

To what extent is CA2 specialized for social compared to non-social odors? We trained mice (n = 5) in the odor-reward association task on a set of four odors (Figure 7A): two novel social odors (urine G and H) and two non-social odors (methyl butyrate and ethyl acetate), from which urine G and methyl butyrate were paired with a reward. Mice were able to learn the task with surprising speed and accuracy (Figure 7B). ROC analysis performed after learning identified cells highly selective to all four odors (Figure 7C). This was accompanied by clearly separated clusters for each odor in principal component space (Figure 7D). Moreover, the social odor clusters versus non-social odors were separated into distinct regions of PCA space, as were the rewarded versus non-rewarded odors. However, in contrast to the purely social-odor trials, we could not identify a clear odor classification schema along any of the principal axes.

**Figure 7.**
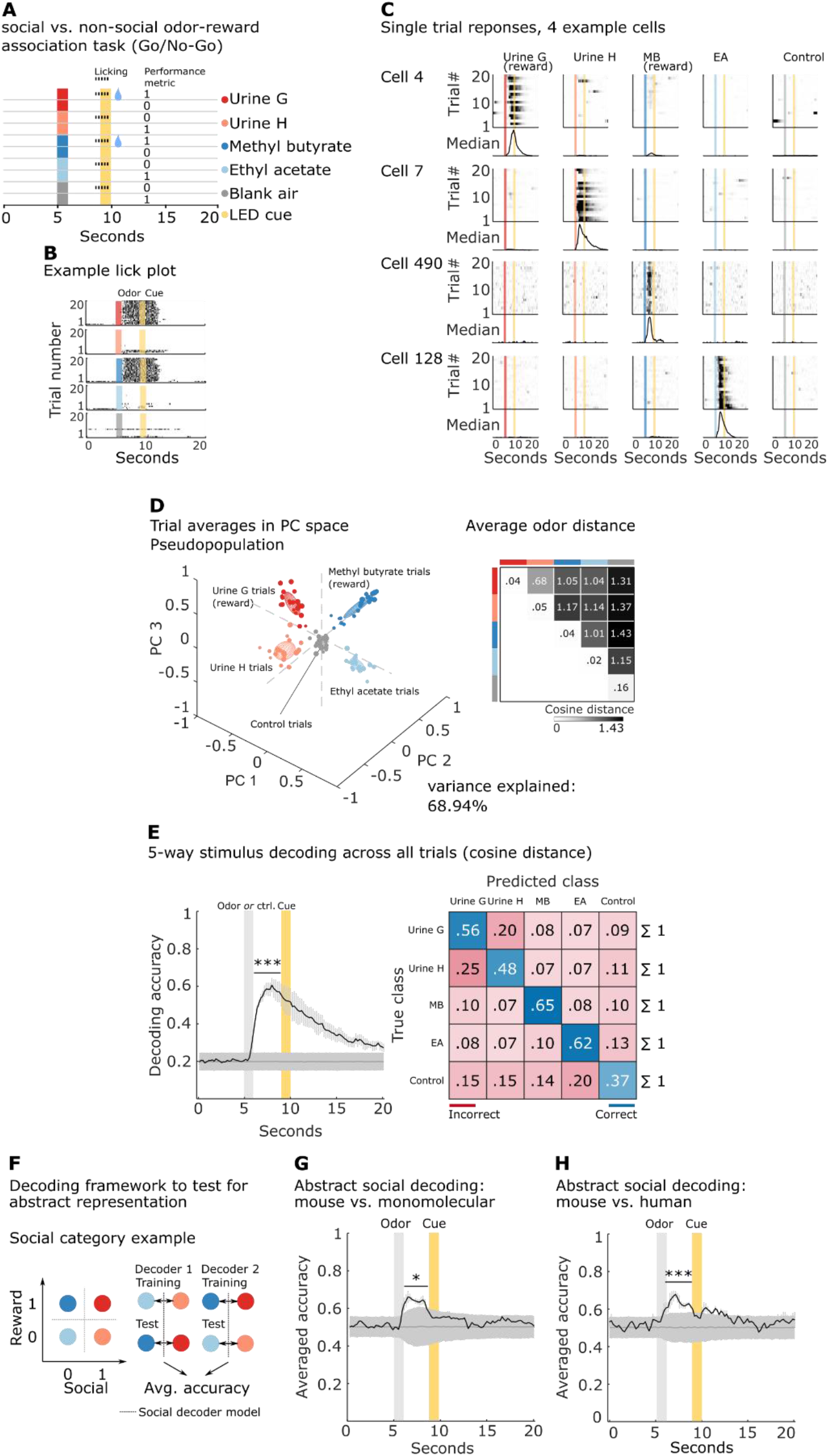
CA2 forms an abstract representation of social versus non-social odors. (A) Schematic of odor-reward association task using two social and two non-social odors, one of each being rewarded, and outcome metric. (B) Example lick behavior of an animal performing above criterion level (>80% correct). Color code as in (A). (C) Single-trial responses of four example cells, each tuned specifically to one of the four odors. Trials are re-sorted and grouped by stimulus condition. (D) Left, PCA representation of CA2 pseudo-population trial responses (533 cells from five animals). The activity was averaged across a three-second window after stimulus presentation (social odors, non-social odors, or blank air control). Right, cosine distance in principal component space (7 principal components, variance explained > 90%). (E) Left, average stimulus classification accuracy of four odors plus blank air (black trace) or chance accuracy (dark grey trace) using the cosine distance of population vectors as a function of time. Results are the average over five animals with 1000 repetitions using random sub-samples of 70 simultaneous recorded cells per animal (total number of cells per animal: 99, 146, 114, 92, 82). Right, confusion matrix summarizing the performance of the classifier. Correct classification is shown in blue along the main diagonal; misclassifications are red-coded off-diagonal values. Darker shades show greater values. (F) Schematic of the decoder test for abstract representation. Left panel, four odor stimuli placed in matrix according two categorical dimensions: a social (circles) versus non- social (squares) odor dimension and a rewarded (darker colors) versus non-rewarded (lighter color) odor dimension. Within-category dichotomies (0/1) lay on opposite sides of a classifier (dashed lines). The classifier was trained to decode one pair of odors across one category dimension (e.g. social versus non-social rewarded odors) and then tested on the remaining pair of odors across the same category dimension (e.g. social versus non-social non-rewarded odors). The classifier was trained and tested across both types of dichotomies for a given dimension and the accuracy was averaged. (G) Decoding accuracy across the social/non-social category dimension as a function of time. The black trace shows the mean decoding accuracy of the two linear decoders (SVM) as outlined in (F), averaged over five mice; mean ± SEM. The first decoder was trained to distinguish between social and non-social non-rewarded odors (urine H and ethyl acetate). The trained model was then asked to classify the two distinct odors of the remaining pair: social versus non-social rewarded odors (urine G and methyl butyrate). The second decoder was trained to distinguish social from non-social rewarded odors and was then tested on the pair of non-rewarded odors. Chance decoding was estimated by decoding with shuffled trial information; *, p < 0.05. (H) Same as in (G) except using a novel set of mouse urine samples I and J and two human urine samples as non-social odors; n = 3, ***, p < 0.001.

We trained and tested a cosine-distance-based classifier for stimulus identity in the full vector space and found highly significant decoding accuracy for all odors time-locked to the odor onset (Figure 7E, left). The confusion matrix of the classifier revealed high true positive rates for all odor classifications, with slightly more errors in distinguishing the two social odors from one another (Figure 7E, right).

### CA2 encodes abstract representations of social versus non-social information

The PCA results suggest that CA2 may represent social versus non-social odors in distinct, abstract categories, as recently described for variables like context, value, and action in primate neocortex and hippocampus (Bernardi et al., 2020). We, therefore, adopted the same analytical approach to determine whether CA2 differentially encodes social versus non-social odors and of rewarded versus non-rewarded odors in a neuronal firing space geometry that permits abstraction. We first trained an SVM binary linear decoder to distinguish the social from the non-social odor using only data from the trials using non-rewarded odors. We then asked whether the trained model could distinguish the social versus non-social rewarded odors, a difficult task due to the difference in valence across conditions and the fact that the trained classifier is entirely naïve to the test data set (Figure 7F). As a result, successful decoding is only possible if there is a specific geometry in the representations such that the same SVM hyperplane that separates the neural responses to the pair of social and non-social non-rewarded odors will also separate the neural responses to the different pair of social and non-social rewarded odors. Moreover, value should be encoded in a subspace that is approximately orthogonal to these two. To account for any asymmetry in the data, we trained a second decoder on the rewarded odor pair and tested it on the non-rewarded odor pair, and calculated the average of both results. The mean performance of this cross-condition classifier showed a decoding accuracy that was significantly greater than chance (Figure 7G). This is not the case when we looked for an abstract representation of reward (Figure S6B). Importantly, both social–non-social and reward–non-reward odor categories can be decoded when a classifier is trained on responses to all four odors for both training and testing (pairwise difference signal minus chance accuracy: 0.34 mean ± 0.07 SEM; p_empirical_ < 0.001, 0.23 mean ± 0.07 SEM; p_empirical_ < 0.001; Figure S6), excluding the trivial explanation that information for only one category is encoded in CA2.

One caveat with the above results is that the abstract decoding could be based on complexity, with social odors being more complex than the monomolecular odorant of the non-social odors. To explore this possibility, we trained three separate animals on a four-odor task as before, using two novel mouse urine samples and human urine samples from two individuals. Human urine is a neutral stimulus for mice (Rivard et al., 2014), with the high complexity of a biological sample. In agreement with the results using monomolecular odors, we found significant abstract decoding in the mouse versus human social category (Figure 7H; pairwise difference signal minus chance accuracy: 0.13 mean ± 0.05 SEM; p_empirical_ < 0.001). Interestingly, using the two sets of complex odors from different species, reward could now be decoded in an abstract manner (Figure S6C; post-odor window: pairwise difference signal minus chance accuracy: 0.09 mean ± 0.04 SEM; p_empirical_ = 0.007; post-reward window: pairwise difference signal minus chance accuracy: 0.1 mean ± 0.06 SEM; p_empirical_ = 0.01). The consistent significant decoding accuracy of the cross-category decoder using both, monomolecular odors and human urine samples, indicates that the abstraction does indeed represent a distinct classification of social (mouse) versus both simple and complex non-social odors.

## Discussion

Our results show for the first time that populations of neurons in the brain encode the identity of complex, highly specific social odors that are necessary to permit animals such as rodents to identify and remember other individuals of their species to form complex social structures. We focused on the CA2 region of the mouse hippocampus because of its well-characterized role in the storage, consolidation, and recall of social memory (Hitti and Siegelbaum, 2014; Meira et al., 2018; Stevenson and Caldwell, 2014). Our finding that CA2 responds to novel social odors in an initial non-specific manner that with experience is replaced by an ability to discriminate between individual social odors provides mechanistic insight into previous electrophysiological results showing that CA2 firing responds both to social novelty (Alexander et al., 2016; Chen et al., 2020; Donegan et al., 2020) and can discriminate social identity (Oliva et al., 2020) during interactions with novel and familiar conspecifics. The activation of a relatively small population of highly selective neurons in CA2 to social odors is in distinction to the finding that spatial information in other regions of hippocampus is mediated by a distributed code of mixed selectivity neurons (Stefanini et al., 2020). Such specific olfactory signals in dorsal CA2, furthermore, are likely to contribute to social engrams previously described in ventral CA1 (Okuyama et al., 2016) through the longitudinal projections dorsal CA2 sends to ventral CA1 that are needed for social memory (Meira et al., 2018).

For social memories to be informative, they need to provide information not only about an animal’s identity but also must contain information about past interactions with a given conspecific, assigning value based on experience. We found that CA2 social representations could be modulated by experience in an associative reward-learning task, leading to an increased discriminability of rewarded and unrewarded social odors. We envision that similar associative learning may enable a given conspecific to be associated with past negative or positive experiences during social encounters. This may underlie the ability of CA2 to regulate social aggression (Leroy et al., 2018).

One intriguing aspect of our findings is that although reward learning increased the fraction of CA2 pyramidal neurons that were selectively activated by a given odor, there was an even larger increase in the fraction of CA2 pyramidal cells that were inhibited by a given social odor. What circuit mechanisms might account for the dual effects of reward learning to increase the populations of excited and inhibited neurons? Although excitatory inputs to CA2 normally show little long-term synaptic plasticity, activation of these inputs can induce long-term depression of feed-forward inhibition (Leroy et al., 2017, 2021; Piskorowski and Chevaleyre, 2013), potentially explaining the fraction of neurons that exhibit increased firing to a given social odor following reward association. A potential mechanism for the increased inhibition is suggested by a recent report describing a powerful direct top-down control of hippocampus by prefrontal cortex, via a specific subset of inhibitory long-range projection neurons. These projections preferentially inhibit vasoactive intestinal polypeptide expressing interneurons in the hippocampus, which result in disinhibition of hippocampal interneurons mediating feed-forward inhibition, leading to a net increase in inhibitory drive onto hippocampal pyramidal cells (Malik et al., 2021). The connection between prefrontal cortex and hippocampus is interesting in light of the finding that prefrontal cortex neurons of mice can form distinct representations of urine pooled from either several male or female conspecifics, although whether prefrontal cortex can encode representations of specific individual social odors remains to be determined (Levy et al., 2019).

What is the nature of the circuit that provides complex social odor information to CA2? In addition to the potential inhibitory input from prefrontal cortex, the major excitatory pathway by which CA2 likely receives social odor information is through the strong direct projections it receives from lateral entorhinal cortex (Chevaleyre and Siegelbaum, 2010; Li et al., 2017; Lopez-Rojas et al., 2021), a brain region that receives strong input directly from the olfactory bulb and indirectly from primary olfactory cortex (Burwell et al., 1995; Leitner et al., 2016) and encodes both individual social sexual odor information as well as changes in novelty (Petrulis et al., 2005). In addition, CA2 receives information about social novelty from its excitatory inputs from the supramammillary nucleus (Chen et al., 2020), although this input targets mainly CA2 inhibitory neurons. Finally, CA2 receives several modulatory inputs, including inputs from neurons in the hypothalamus that release the social neuropeptides vasopressin and oxytocin (Cui et al., 2013; Lin et al., 2018; Raam et al., 2017; Smith et al., 2016; Tirko et al., 2018). An important goal of future studies will be the identification of the specific role these diverse inputs may play in mediating the complex social odor responses identified in this study.

Our finding of abstract social representation in the hippocampus is remarkable in the light of results from piriform cortex, which although upstream of lateral entorhinal cortex and hippocampus lacks a differential representation of innately relevant versus non-relevant stimuli (Iurilli and Datta, 2017). Our results indicate that the emergence of abstract representation occurs further downstream in hippocampus. It will be of interest in the future to determine whether hippocampus inherits these abstract categories from lateral entorhinal cortex, although abstract representation may first emerge in hippocampus based on fMRI studies in humans as they form a map of social space (Tavares et al., 2015). It will also be of interest to test the existence of an abstract representation of social identity at the level of freely behaving animals, e.g., to determine whether the geometry of social representations allows abstract categorization of novel versus familiar animals or males versus females.

Our finding that CA2 social odor representations are unstable in sessions across more than one is consistent with findings that CA2 representations of space are also relatively unstable (Donegan et al., 2020; Mankin et al., 2015). Although there is significant representational drift of odors in the upstream piriform cortex, these representations are stable over several weeks (Schoonover et al., 2021), implying that CA2 (or lateral entorhinal cortex) employ active mechanisms to de-stabilize representations. What advantage could this have? Social behavior is notoriously dynamic. An overly rigid and stable representation might handicap necessary adaptions and prove harmful on a behavioral level. Recent models suggest, based on experimental evidence, that there are computational advantages for the brain to enact a compromise between persistence and flexibility (Mau et al., 2020; Richards and Frankland, 2017; Rule et al., 2019). Indeed, CA2 activity in a mouse model of the human 22q11.2 microdeletion, which is strongly linked to schizophrenia, formed an overly stable representation of space, which was associated with impaired CA2 social firing and a deficit in social memory compared to controls (Donegan et al., 2020). In light of our findings, it will be interesting to determine whether persistent tuning of CA2 may serve as a general pathophysiological mechanism for social memory deficits in other models.

## Conclusion

Our results provide evidence for the representation of social olfactory signals as the likely candidate underlying CA2’s ability to encode and recall social memory. They strongly argue for a role of CA2 beyond novelty recognition and provide direct evidence for a causal role of CA2 to flexibly integrate information like valence in a social representation. The existence of abstract representation speaks for CA2 being a powerful hub in the organization of multidimensional social features into a coherent abstract social map in the hippocampus. Our results and experimental framework provide the basis to specifically expand on questions regarding the relationship between social novelty and familiarity and the flexible representation of a social entity in both health and disease.

## Acknowledgments

We thank Jennifer Bussell and Clay O. Lacefield for their initial help with the olfactometer design; Darcy Peterka for help in setting up the two-photon microscope; Alex H. Williams for discussion on the tensor analysis; Walter Fischler-Ruiz, Daniel Salzman, Stefano Fusi, Arjun V. Masurkar, and Lara Boyle for critical comments on the manuscript; Franziska Auer and the Siegelbaum laboratory for helpful discussions; the Research Instrumentation Core at Zuckerman Institute. This work was supported by a NARSAD Brain and Behavior Research Young Investigator Award (to S.I.H.), grants MH-104602 and MH-106629 from the National Institute of Mental Health (NIMH), and a grant from the Zegar Family Foundation (to S.A.S.).

## Author contributions

S.I.H. and S.A.S. conceived of the project and wrote the manuscript. S.I.H. designed the experimental setup and performed all experiments except S.B. performed the optogenetics experiments, immunohistochemistry, and contributed recordings to imaging experiments using human urine samples. S.I.H. wrote software and analyzed the data.

## Experimental model and subject details

All mouse procedures were performed in accordance with the regulations of the Columbia University IACUC. We used heterozygous sexually naïve Amigo2-Cre (Hitti and Siegelbaum, 2014), 3-6 month old female mice except for experiments including human urine samples, where we used a mixed group of male and females (no qualitative difference in behavioral performance between genders was apparent in this group), on the C57Bl/6J background (The Jackson Laboratory. All mice were maintained on a 12-h light-dark cycle with ad libitum access to food. For behavioral training and imaging, mice were kept on a water schedule, where they received 1 ml in total per day. Mouse health was monitored daily and we gave additional water if their weight fell below 80% of their pre-schedule weight. Mice were housed in groups of 2-4.

## Methods detail

### Stereotaxic viral delivery

For imaging experiments, mice were injected with a bicistronic Cre-dependent AAV2/1 expressing the genetically encoded activity indicator GCaMP6s and nuclear tdTomato, with the latter serving as a structural marker to improve motion correction AAV-EF1a-DIO-GCaMP6s- P2A-nls-dTomato was a gift from Jonathan Ting (Addgene viral prep #51082-AAV1; http://n2t.net/addgene: 51082; RRID: Addgene_51082). Injections were targeted unilaterally to the left dorsal CA2 region using Allen Brain Atlas coordinates: anterior-posterior (AP): -1.7, medial-lateral (ML): 2.0, from bregma; dorsal-ventral (DV): -1.3 from pia mater. ∼60 nl in saline diluted virus (titer: 1*10^12^ gc/ml) were delivered at 1 nl s^−1^ using pressure injection. A cranial window was implanted 24h after injections.

In silencing experiments, AAV2/5-EF1a-DIO-hChR2-eYFP (titer: 4.0*10^12^ gc/ml) or AAV2/2- EF1a-DIO-mCherry (titer: 4.4*10^12^ gc/ml was bilaterally injected (200 nl per site) to express the inhibitory opsin eArch3.0 or a fluorescent control selectively in CA2. Right after injections, optogenetic fibers were implanted.

### Surgical implants

Mice were implanted with an imaging window (diameter 3 mm, height 1.8 mm) over the left dorsal hippocampus and a custom stainless-steel headpost to allow for head fixation during imaging. Imaging cannulas were constructed by adhering (Norland optical adhesive) a 3 mm glass coverslip (Warner instruments) to the steel cannula (Ziggy’s tubes and wires). For the implant, mice were anesthetized (isoflurane) and provided with anti-inflammatory (dexamethasone 2 mg/kg, s.c.) and analgesic (buprenorphine SR, 1mg/kg, s.c.) treatment. A single 3 mm craniotomy, centered at the site of injection, was performed using a trephine drill. Cortex was irrigated using ice cold sterile saline supplemented with 3 mM MgCl2 and 1 mM CaCl2 and carefully removed through vacuum aspiration to allow visual access to the hippocampus. The prepared optic cannula was inserted at a 15° angle and secured with Vetbond (3M). Both cannula and head-post were then permanently adhered to the scull using dental cement (Metabond, Parkell). Mice recovered in their home cage, and behavioral training started 2 weeks after surgery.

Optical fiber assemblies (200 um core, 0.37 NA, 3 mm, RWD Life Science Inc.) were implanted at the site of injections and placed on top of the alveus. Fibers were permanently fixed using dental cement.

### Immunohistochemstry

Posthoc immunostaining of brain sections were performed as previously reported (Leroy et al., 2018). Briefly, mice were perfused with chilled 4% PFA in PBS. 60 μm coronal sections were prepared and area CA2 was counterstained using rabbit anti-PCP4 (1:400, Sigma-Aldrich, #HPA005792).

### Odor selection

Urine samples from individual male mice (1-3 ml, form∼ 3 month old animals) and humane urine samples (male, 20-30 years old) were ordered (BioIVT) and diluted in water (1:50). Ethyl acetate and methyl butyrate (Sigma) were diluted in mineral oil (1:200 and 1:2000, respectively).

### Behavioral setup

A custom 8-channel olfactometer was built and controlled using custom written Matlab (Mathworks, R2020b) code. A constant stream of 1 l/min medical air and a second, independent carrier stream (0.5 l/min) were combined at a nose cone, which was placed 0.5 cm in front of the head-fixed animal’s nose. The carrier line contained either air that was guided over individual reservoir bottles containing the respective odors or simple water as blank control. A microcontroller (Arduino) controlled valve array (The Lee Company) switched precisely between blank and odor bottles, which resulted in a total air stream of 1.5 l/min across a session. The water-delivery system consisted of a valve-controlled, gravity driven water supply that was connected to a 22G gavage needle, which was held in place ∼3mm in front of the animal’s mouth. A capacitance sensor (Sparkfun, #AT42QT1010) was soldered onto the gavage needle to register lick behavior. The sampling rate was set to 20 Hz. A white LED, which indicated the cue window, was placed ∼4 cm in front and slightly above the animal’s head. To synchronize the behavioral setup with the two-photon image acquisition, both the microscope and the olfactometer were set to send continuously TTL pulses to a master data acquisition board (National Instruments, #USB-6001), which sampled at 500 Hz.

### Behavioral training

Throughout an experiment, mice were head fixed in a custom-printed enclosure which prevented larger movements but provided enough space for animals to adjust their posture. Mice were initially habituated to be head fixed by just placing them repeatedly in the enclosure until they showed no obvious sign of discomfort (e.g., agitation). In all following training phases, the amount of water received during training was registered and the difference to the 1 ml/day water schedule was provided to the animal in a dish placed in a separate cage after training. In the initial ‘shaping’ phase, mice were trained to associate a light cue with water delivery (4 μl) through the lick port. The protocol for this phase consisted of 20, 4-second long unconditional trials, where a 5-s baseline was followed by a 1-second LED cue. After the cue a water droplet was delivered. The 20 unconditional trials were followed by a maximum of 80 conditional trials, where the 5-seond baseline was followed by the LED cue, however, now the animal had to actively engage by licking at the lick spout to immediately trigger a water reward. If the mouse did not lick within 4 seconds, during which the LED was continuously on, the trial was considered a failed trial and a new trial was initiated. A maximum of 200 attempts was given on a single day to reach 80 successful trials. Once the animal consistently reached 80 success trial in 80 attempts, the animal moved to the ‘cue-training’ phase. Here a single trial lasted 10 seconds. An 8-second long baseline was followed by a 1-second long LED cue, in which the animal had to actively lick at the lick spout. If it did, it received a water reward right after the cue window and a new trial was initiated. If it failed, no punishment was imposed upon the animal and the new trial was initiated. The animal had to complete 100 trials per session and the performance was monitored as the fraction of successful trials over all trials. Once the animal performed > 90% correct trials, the animal moved to the actual ‘odor-discrimination’ phase. In this phase a single trial lasted 20 seconds: a 5-second baseline, a 1-second odor/blank presentation, a 3-second delay, a 1-second cue-window, and a 10-secodn inter-trial-interval.

Depending on the paradigm (2 or 4 odors discrimination), 1 or 2 odors were paired with a reward, and the mouse had to actively lick at the spout during the cue-window after those odors were presented, but not to the others. Licking after the correct odors and withholding to the non-rewarded odors or blank trials, was counted as a correct trial and the performance measure was the fraction of correct trials over all trials. Mice underwent 90 trials per session in the 2-odor task and 100 trials in the 4-odor task, which kept the total session time > 40 min., an empirical value in which mice showed consistent engagement in the task. To enhance engagement of the animal in the task, we started each session in the discrimination phase by 3 consecutive presentation of the rewarded odor(s), where we kept the trial structure, however we presented a reward after the cue-window unconditionally of the animal’s licking behavior.

During optogenetic silencing we inhibited CA2 pyramidal neurons by activating inhibitory archaerhodopsin expressing in CA2 of Amigo2-Cre mice by shining light (10 mW laser output, 532 nm) 1 seconds before odor onset, lasting 2 seconds into the inter-trial-interval to inhibit both, encoding and recall processes.

### Two-photon calcium imaging

Imaging was performed using a galvo-galvo (Cambridge Technologies) based 2-channel Scientifica scope. Acquisition was performed with a large working distance Olympus 20x air lens (LCPL20XIR, 0.45 NA, 8.93 - 8.18 mm WD). A Mai Tai laser (Spectra Physics) tuned to 920 nm delivered 50-100 mW of excitation power at the front end of the lens. Emitted fluorescence was detected using a GaAsP detector on the green channel and a bialkali on the red one (MDU-PMT-50-00 Small B.A GaAsP and 2PIMS-PMT-40, respectively; Scientifica/Hamamatsu).

Scanimage 5 was used for hardware control and data acquisition.

For each animal we acquired a z-stack spanning 50 μm centered at the imaging plane before the first recording session. This allowed to take advantage of Scanimage’s correlation based online ‘motion-estimator’ module and we identified the exact field-of-view on each recording day. In addition, it allowed to monitor x-, y-, and z-drift during recordings, which we occasionally corrected manually during a recording session if it exceeded the threshold of 5 μm. A field-of-view covered ∼200 by 200 μm and we scanned at 14 Hz. Both green and red signal were recorded, with the latter serving as input signal for software based offline image-alignment and motion correction.

### Signal extraction and longitudinal registration

Detection of regions of interest (ROIs), segmentation, and extraction of fluorescence signal was performed using the Suite2p software (Pachitariu et al., 2017). This package implements single day image registration and fluorescence source and neuropil detection from spatially overlapping ROIs. All identified ROIs were confirmed by manual inspection. Suite2p also provides a principal component analysis-based motion estimator module (Stringer and Pachitariu, 2019), which allows to qualitatively detect even subtle remaining drifts after offline motion correction. We only included animals with stable window implants where we could not detect motion drifts after correction. ROIs where excluded where we detected nuclear expression of GCaMP, a sign of toxicity caused by overexpression.

For day-to-day longitudinal cell registration we used a pipeline reported earlier (Sheintuch et al., 2017) and used the probabilistic model (psame threshold = 0.5) based on spatial correlation of the identified cell masks.

### Data processing

Extracted fluorescent traces were neuropil subtracted (scaling factor for neuropil: 0.7) and slow baseline trends were corrected by subtracting the 8^th^ percentile within a 30 second window as reported earlier (Dombeck et al., 2010). Detrended traces were scored by their median absolute deviation. To further correct for differences in expression level we followed the framework described earlier (Mackevicius et al., 2019) and we re-normalized signals by dividing each neuron’s signal by the sum of its maximum value during a session and the 95^th^ percentile of the signal across all neurons. In this way, we reduced major differences between strongly and weaker expressing cells, but still retained priority for larger transients. To align signals from the microscope with behavioral data, which were sampled at different frequencies (14 Hz and 20 Hz respectively), we aligned signals using nearest-neighbor-interpolation and re-sampled at 10 Hz.

## Data analysis

Data analysis was performed using custom written code in Matlab and Jasp (v0.14.1) for repeated-measure ANOVA analysis.

### ROC analysis

To detect significant activation of a cell above baseline we used receiver-operator-characteristic (ROC) analysis. We generated 2 activity distributions, the average fluorescent activity for each trial of a respective cell-odor pair in a 3-second time window before (t = 1-4 s during a given trial) and after (t = 6-9 s) odor presentation. The resulting 2 activity distributions were subjected to ROC analysis and we computed the area-under-the-receiver-operator-curve statistic (AUC) (see also Figure S1A) to derive the selectivity-index as: 2 * (AUC - 0.5). The index is bound between -1 and 1. A value of 0 indicates no bias to either baseline or post-odor presentation. A value below 0 indicates higher activity during baseline, whereas values larger than 0 a higher activity after odor presentation. To determine the significance, we sampled under the null-hypothesis (no difference between the distributions) and re-calculated a selectivity index. We repeated this 1000 times, which provided for each cell-odor pair a selectivity-index distribution. A significant response was defined whenever the experimental AUC value was either below the 2.5^th^ percentile or above the 97.5^th^ percentile of the shuffled distribution. To determine the selectivity of responses to the two odors, rather than odor versus baseline, we used the post-odor activity distributions for both odors for each cell and subjected these to the ROC analysis.

### PCA analysis

For principal component analysis (PCA) we first averaged each cell’s fluorescence response signal within the respective trial class. We focused only on values during the 3 s after odor presentation (post-odor period). We constructed a data matrix with individual cells in columns and concatenated the averaged median-absolute-deviation scored normalized response in the post-odor period for each trial class along rows (e.g., 3 s period sampled at 10 Hz resulted in 30 data points for a trial type; with 2 odors and control blank air trials this resulted in 90 data points for each cell). We then calculated the covariance matrix for correlations in mean neural activity among the different cells and used PCA analysis to identify the eigenvectors of the covariance matrix (principal components). To determine the contribution of each principal component to a given trial we projected the matrix containing the original fluorescence responses of a given trial for all neurons onto each component. We then averaged the projected values within each trial and plotted these values for the first 2 or 3 principal components.

### Tensor decomposition

For tensor decomposition analysis (TCA) we used the Tensor Toolbox for Matlab (Brett Bader, Tamara G. Kolda et al., version 3.2.1, 2021). We formed a 3D input matrix with the dimensions: *cells* x *time* x *trials*. We used the tensor() function to create a tensor class from this matrix and then ran the cp_opt() function for the decomposition. We used random initialization, set a lower threshold of 0, to ensure non-negativity and fit a 12-component model with otherwise standard settings.

### Decoder analysis

For binary decoders, we implemented a Support Vector Machine (SVM) performing C-Support Vector Classification with a linear kernel using the LIBSVM Matlab implementation (Chang and Lin, 2011). We used 5-fold cross-validation throughout our decoding analysis. The regularization hyperparameter *C* was selected using a random search on a separate validation set. We chose a less permissive value of 50 for all experiments. To decode across a single-trial time-course, we averaged fluorescent data within half overlapping bins using a bin size of 250 ms and trained and tested a separate decoder on each bin. Chance decoding was accomplished by randomly shuffling trial identity. We used random subsampling from the total pool of cells to ensure consistency in dimensionality when comparing across decoders. Results were calculated as average accuracy across folds and subsamples. The cross-validated subsampling distribution represents the statistical sampling distribution and the estimate for the standard error of the mean is therefore simply given by the standard deviation of this distribution. Significance above chance was determined by first averaging for each subsample across the post-odor period (6-9 seconds) and subtracting the chance accuracy from the signal accuracy. The corrected empirical p-value (North et al., 2002) is then given by: (*r* + 1)/(*n* + 1), with *r* being the number of values from the distribution that are smaller than 0 and *n* the total number of subsamples.

When we trained and tested decoders using individual animals rather than decoding from pseudo-populations, we first sub-sampled within animal and pooled the subsamples to calculate the empirical p-value (e.g., 6 animals, with 1000 subsamples each, resulted in a total distribution of 6000 values).

For the cross-category decoder that tests for abstract representations, we trained and tested in 2 opposite directions: in each subsample, we first trained decoder-models across time to classify the non-rewarded social and non-social odor and tested the trained models on the rewarded odor. This way, the trained model is naïve with respect to the testing data. In the second direction we trained on the rewarded odors and tested on the non-rewarded odors. This bidirectional training/testing accounts for asymmetry in the data. We ran 1000 subsamples per animal and in the end pooled subsamples to estimate the average accuracy and significance as described above.

When we had more than 2 classes to predict, we used a decoding approach based on the cosine distance of the population vectors. We first binned the data using 250 ms bins. For each bin, using 5-fold cross validation, we trained a model by calculating the average population vector for the responses to each odor and the blank air in *n*-dimensional neuron-activity space (*n* = number of neurons; e.g., training in the 2 odors plus 1 blank paradigm resulted in 3 average population vectors) in the training set. We then calculated for each trial in the test set its cosine-distance to the respective average population vectors. The class with the smallest distance became the predicted odor for this trial. We used subsampling with 1000 random samples and calculated the empirical p-value as described above.

### Binomial test

We tested the conditional probabilities across sessions in multiday recordings for four selectivity categories (gained, lost, retained, or reversed selectivity across days; Figure S4). To determine whether the empirical values deviated significantly from the expected values in the case of independent probabilities, we used a two-sided Binomial test. We used the Matlab implementation of Matthew Nelson (Matthew Nelson (2021), myBinomTest(), MATLAB Central File Exchange. Retrieved July 29, 2021.). We first calculated the independent probabilities; for example, for cells that lost their odor selectivity between sessions, this is given by the product of the fraction of cells that were selective on a previous session and the fraction of non-selective in the following session. The conditional probability is the fraction of cells that lost selectivity in the pool of cells identified in both sessions.

### Fitting

To characterize the behavioral learning curve for odor association with a water reward, we used Matlab’s fit() and coeffvalues() functions to fit a 3-parameter logistic model to the data: *a* + (1 − *a*)/(1 + exp(*b* ∗ (*x* − *c*))), with *a* being the maximum/plateau value, *b* the slope, and *c* the midpoint/inflection point. In cases where we tested for significance, we used Monte Carlo random sampling testing. We randomly assigned values from the treatment and control group, fitted the model, and saved the resulting coefficients. We repeated this 5000 times, which resulted in a distribution of coefficients, and then calculated the one-sided, corrected empirical p-value by: (*r* + 1)/(*n* + 1), with *r* being the number of values from the shuffled distribution that are either smaller or larger (depending on the hypothesis) than the observed value and *n* the total number of repetitions, in this case 5000.

---

**Figure S1.**
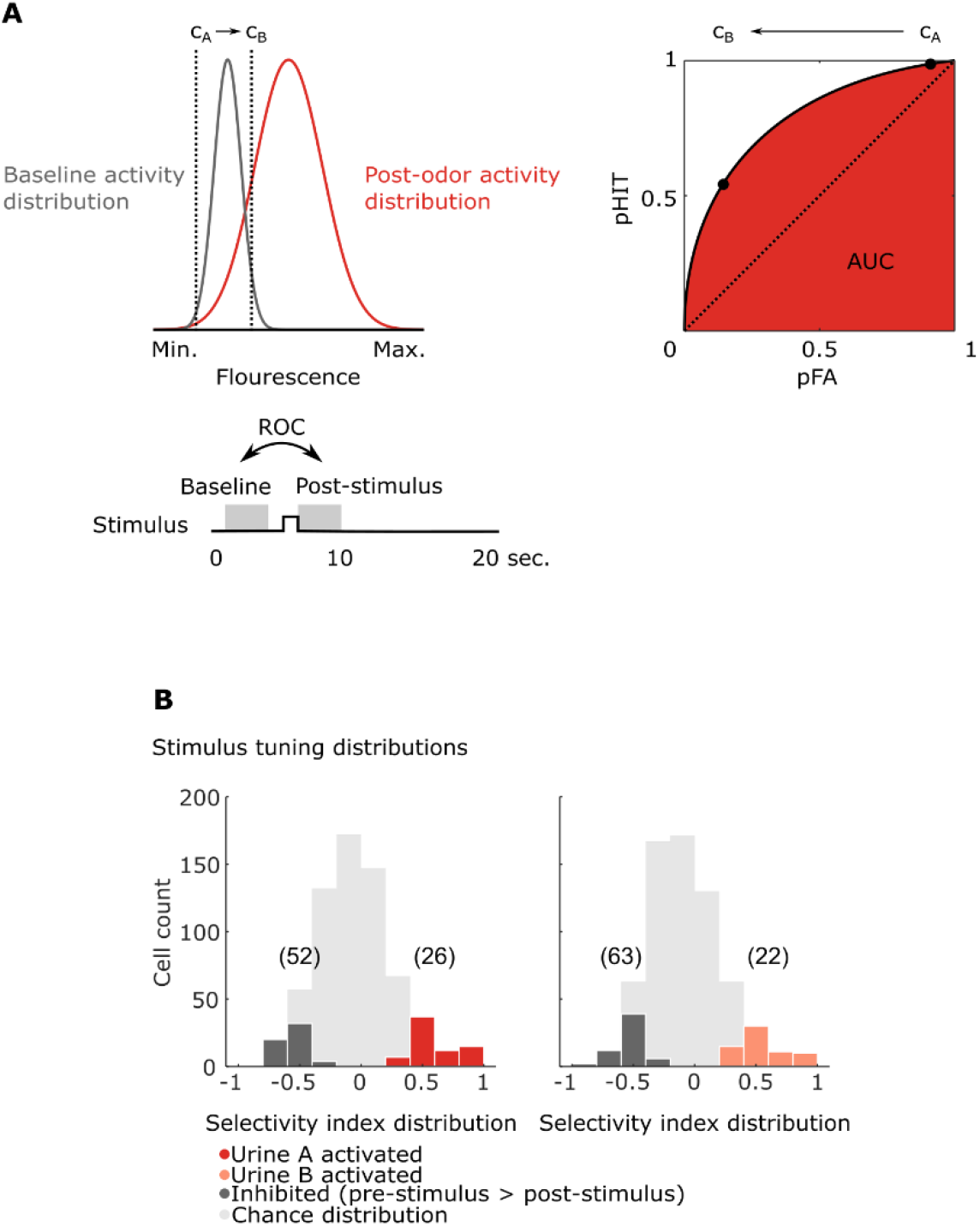
Tuning distribution of CA2 pyramidal cells after passive social odor presentation. (A) Schematic of the Receiver-Operating Characteristic Analysis (ROC). Left, ROC uses a varying threshold *c* (cA and cB) to estimate the separability of two distributions (here fictive Gaussian distributions, modeling average trial activity values under baseline (grey) or post-odor (red) condition). For each value of *c,* the fraction/probability of the distributions to the right of it is registered (in this case, pHIT for the odor distribution and pFA for the baseline distribution). The resulting pairs of probability values for each *c* are plotted against one another and result in a ROC curve. The area under the curve (AUC) represents the probability that a neuron’s response will be ranked higher than a response obtained under baseline activity. The diagonal isoline (dashed line) represents indifference between odor and baseline. (B) Distribution of selectivity indices from data in Figure 1I. Left: the distributions of cells, whose ROC-based selectivity index differs significantly from chance for urine A (dark red) or cells that responded to urine A with decreased firing relative to baseline (dark gray), are overlaid over the non-significantly tuned fraction. Right: same as left but for social odor B; significance was determined by re-sampling under the null-hypothesis; p <, 0.05; 659 cells, 7 animals.

**Figure S2.**
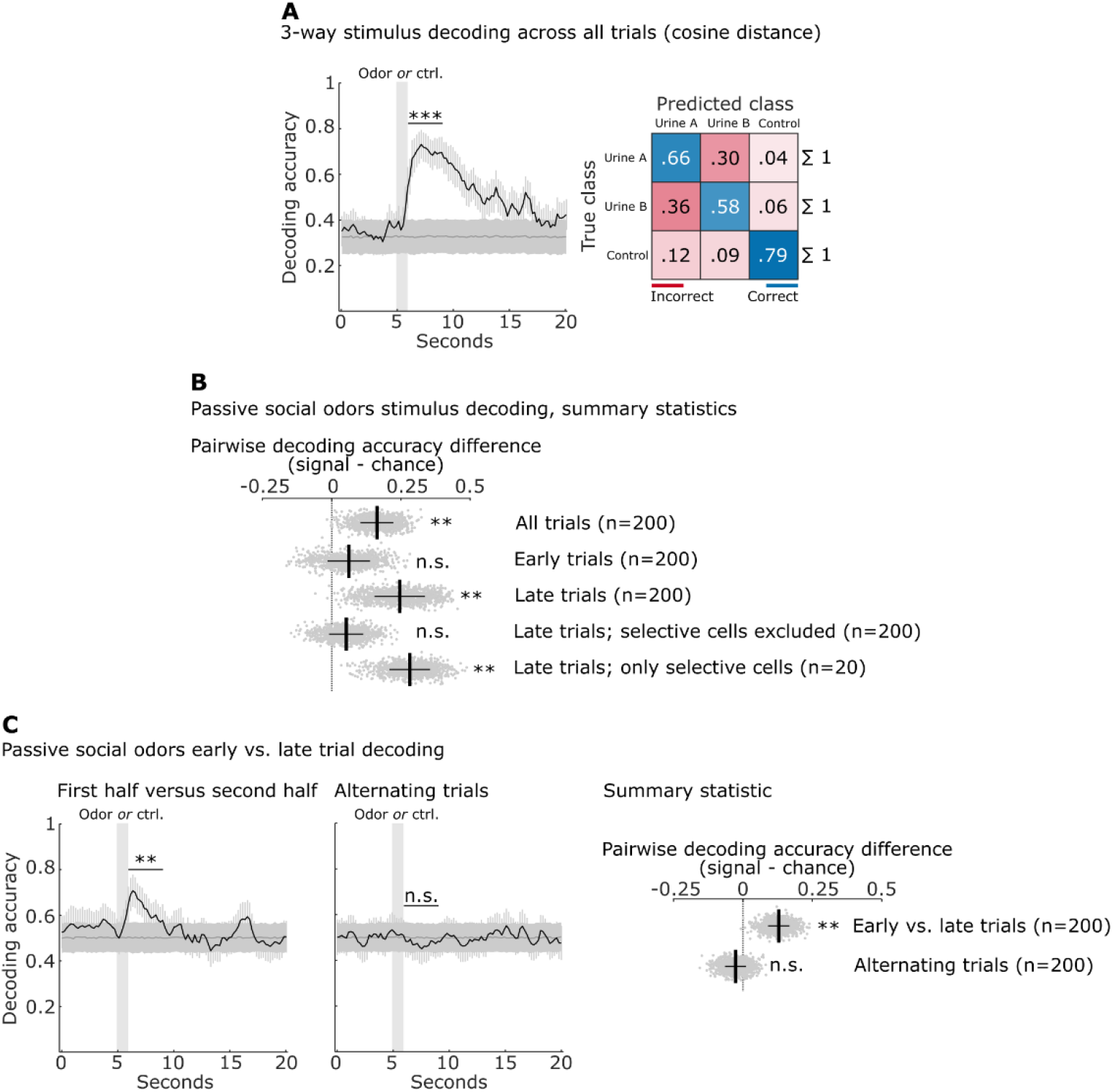
Stimulus and trial decoding in CA2 during passive odor presentation. (A) Left, average stimulus classification accuracy (black trace) or chance accuracy (dark grey trace) using a cosine distance classifier to detect odor identity or blank air control as a function of time. 1000 repetitions using random sub-samples of size 200 from the pseudo-population constructed from the data of seven animals. Right, confusion matrix summarizing the performance of the classifier. Correct classification is shown in blue along the main diagonal; misclassifications are red-coded off-diagonal values. ***, p < 0.001. (B) Summary statistic of stimulus decoding. The distribution of pairwise sub-sampled differences of averaged binary decoding accuracy (SVM) over the three-second window following odor presentation (signal minus chance accuracy; 1000 subsamples, light grey dots). Overlaid is the average (thick black vertical line) and SEM (thin black horizontal line). Each row represents a different decoding condition; empirical p-value were determined to estimate significance (see methods for explanation); **, p < 0.01; n.s., non-significant. (C) Decoding of trial number within a session. First panel: a decoder was trained to classify whether a trial occurred in the first or the second half of a session. Non-selective novelty signals in CA2 allow for significant decoding time-locked to the odor onset and irrespective of odor identity. Accuracy of a decoder trained to distinguish between alternating trials was not above chance (second panel). Color code as in (A); summary statistic, see (B) for explanation; **, p < 0.01; n.s., non-significant.

**Figure S3.**
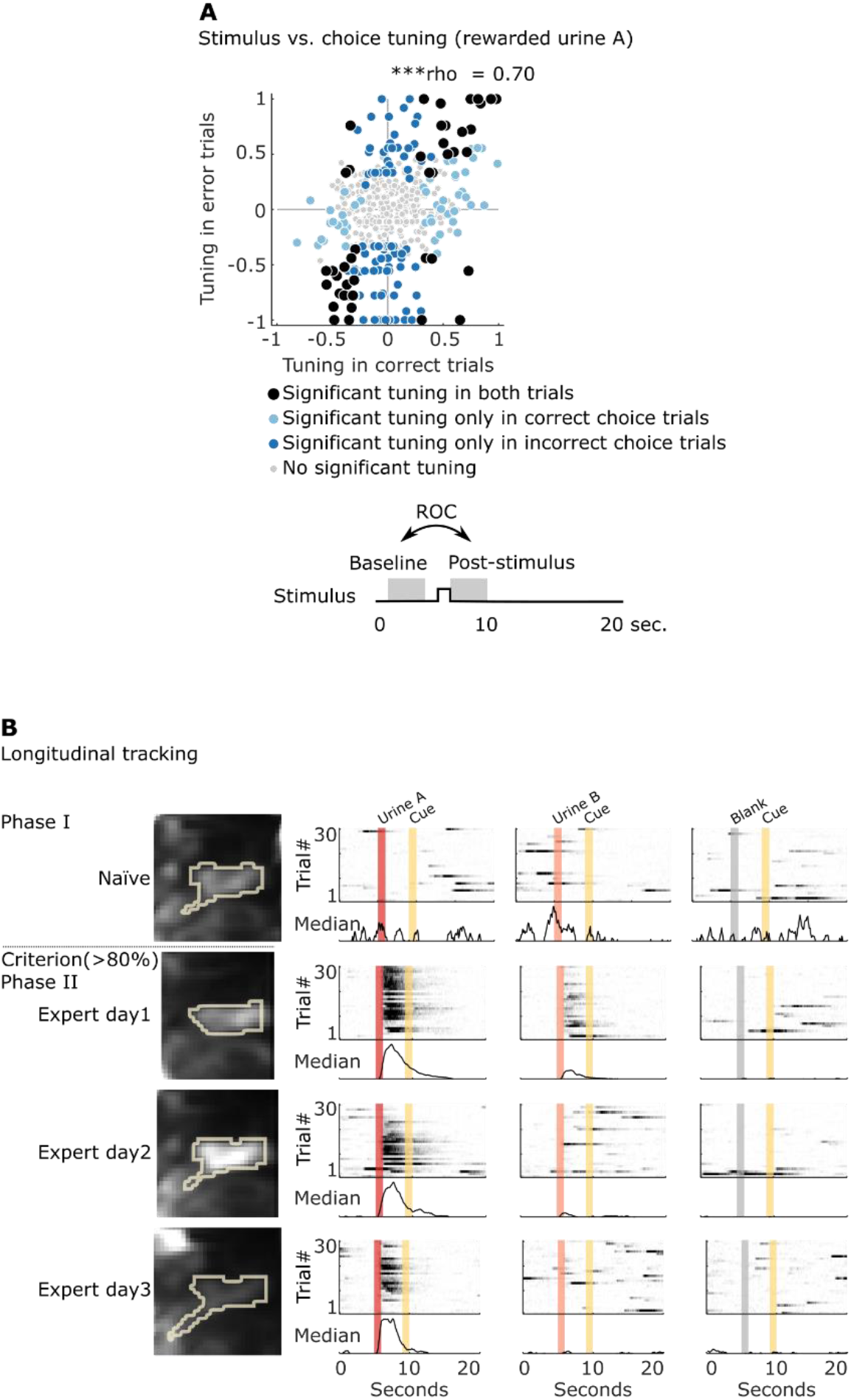
Stimulus versus choice tuning and longitudinal tracking. (A) Shown are the AUC-based selectivity indices (ROC, baseline versus post-stimulus) for all cells pooled across animals (n = 6) on the first day of the associative reward-learning. Indices of cells in error trials are plotted as a function of their indices in correct trials. The positive correlation (Spearman rho) of cells that are significantly tuned in both error and correct trials indicates stimulus tuning rather than behavioral choice (lick vs. no-lick). ***, p < 0.001. (B) Longitudinal tracking. We tracked responses of individual cells to odors and blank air across sessions (Sheintuch et al., 2017). An example cell with its identified region-of-interest mask is plotted across sessions (left). Its corresponding single trial activity in response to indicated stimuli is shown in raster plots on the right.

**Figure S4.**
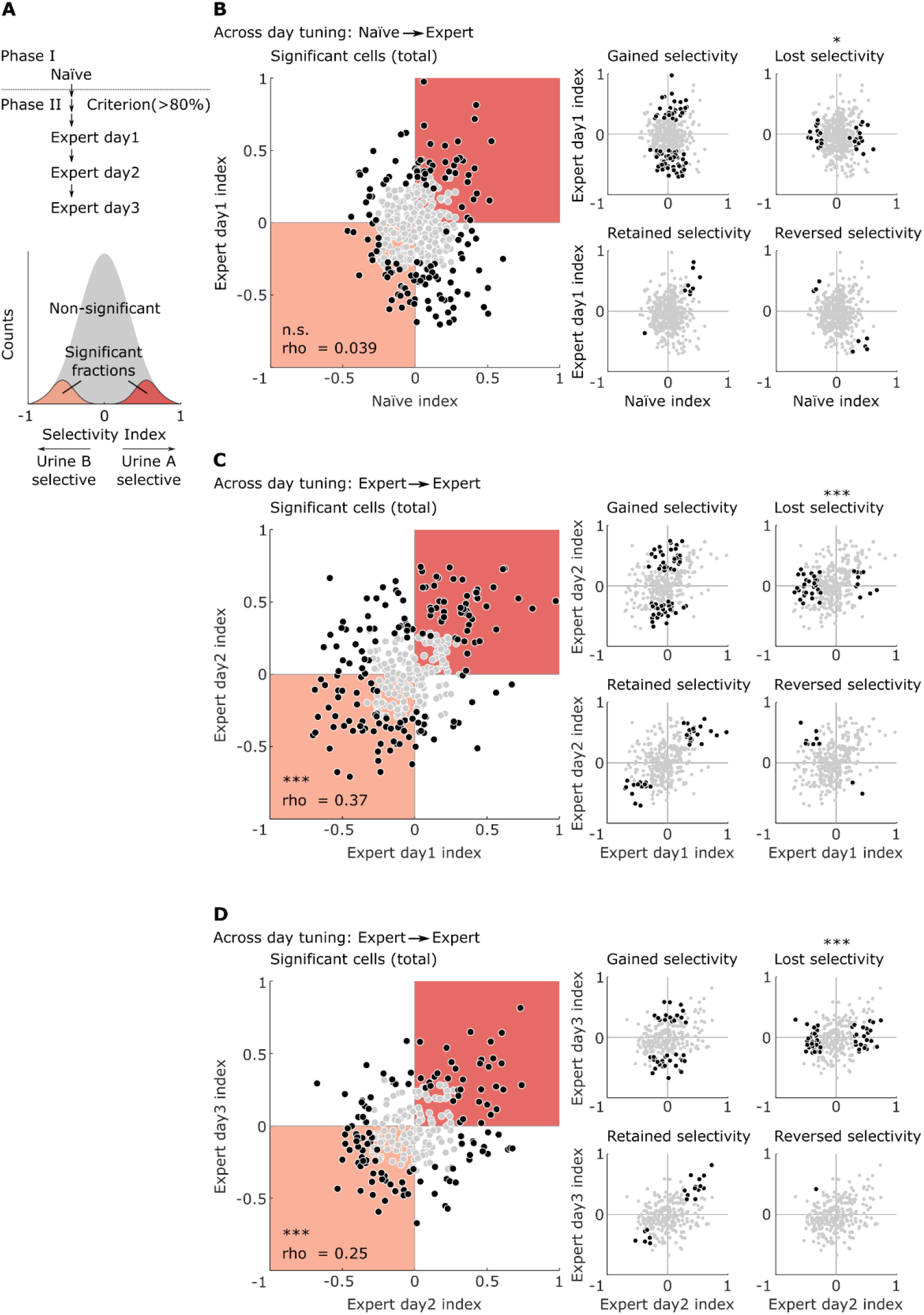
Single cell tuning across days during learning of social odor-reward associations. (A) Schematic of the ROC analysis across days. Selectivity indices were based on the comparison between the trial activity vectors of urine A versus urine B, rather than odor versus baseline, using the ROC framework. Data was formatted in a way that urine A selective cells always had a positive index (bound between > 0 and 1) and urine B selective cells a negative index (bound between -1 and < 0). Cells with a selectivity index of zero had no selectivity bias. The significance level of p < 0.05 was determined using shuffled distributions. (B) – (D) Selectivity indices plotted across days/sessions. The same cells were first identified across two or more sessions and pooled across animals (n= 6, 5, 4 for B-D, respectively). Selectivity indices of cells from one session were plotted as a function of their indices in the previous session. Black points were selective in at least one of the two sessions. The upper-right and lower-left quadrants represent consistent selectivity for urine A and urine B, respectively. From the pool of selective neurons, we could extract four classes: cells that gained selectivity (from B to D: 88, 73, 45), lost selectivity (31, 40, 59), retained their selectivity (99, 42, 19), or reversed their selectivity (9, 10, 1) between successive sessions. The number of total cells identified was 396, 329, and 258. Spearman rank correlation was used to calculate rho, a binomial test (see methods for explanation) was used to test significance of the conditional probabilities; n.s., non- significant, ***, p < 0.001.

**Figure S5.**
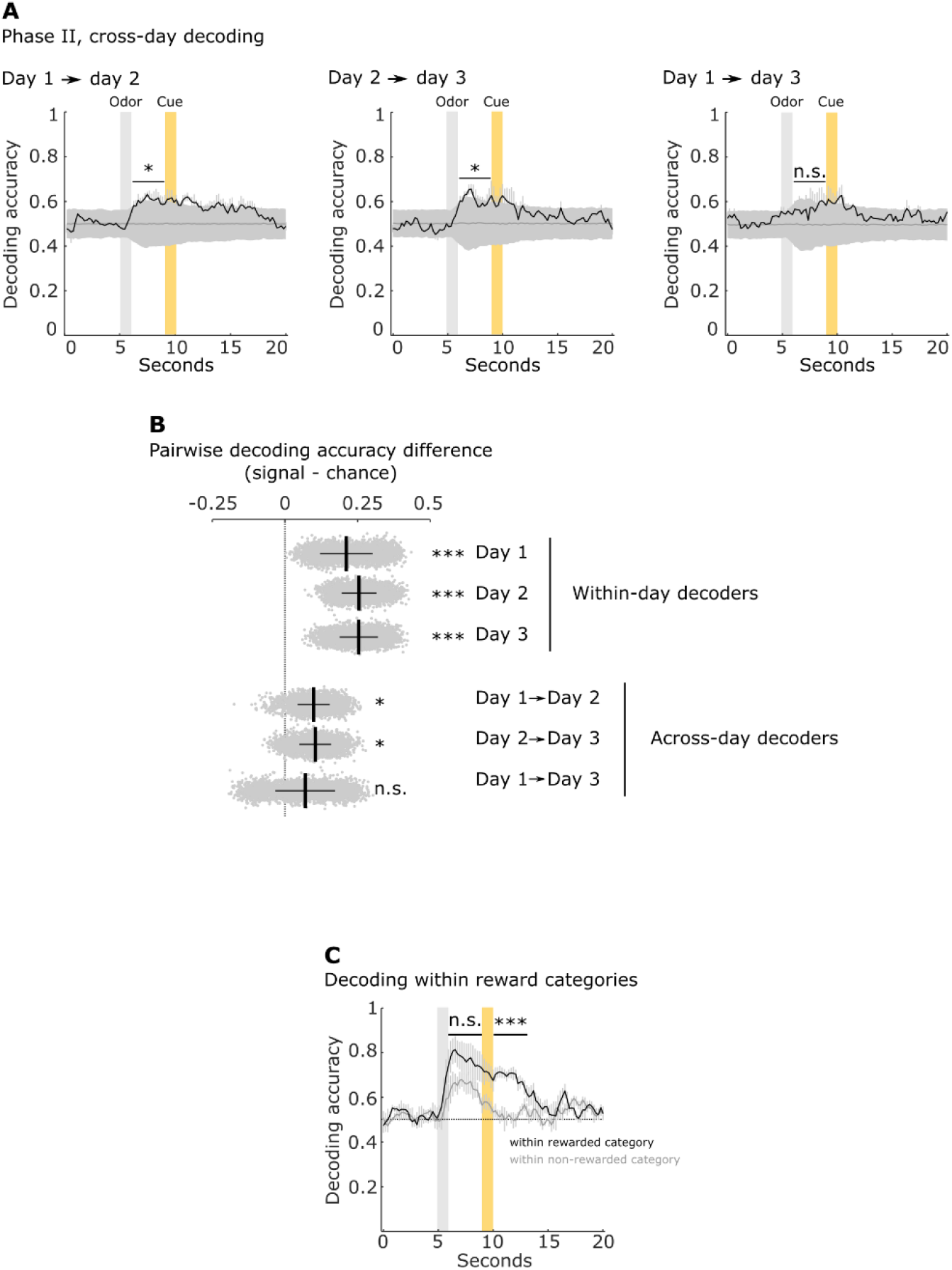
Cross day decoding and decoding within and reward categories. (A) Cross-day decoding in Phase II. Average decoding accuracy of the SVM binary linear decoder trained on one session and tested on an ensuing session using the same set of cells identified over all sessions (70 cells, 1,000 samples per mouse; n = 5). Significance is determined in the three second delay window after odor presentation by calculating an empirical p-value using the pairwise difference distribution between signal and chance distributions; n.s., non-significant; *, p < 0.05. (B) Summary statistic comparing within and across day decoding in Phase II. Description as in Figure S2B. (C) SVM decoding accuracy of two social odors with same reward contingency in the four social odor task. Black trace, decoding of the rewarded odors. Grey trace, decoding of the unrewarded odors. The decoding accuracy for rewarded and non-rewarded odors did not differ significantly during 3-s delay period after odor presentation prior to reward (n.s., non-significant; p = 0.3; n = 3 animals). After reward delivery the decoding accuracy for rewarded odors was significantly greater than that for non-rewarded odors (***, p < 0.001). P-values calculated using bootstrap permutation test.

**Figure S6.**
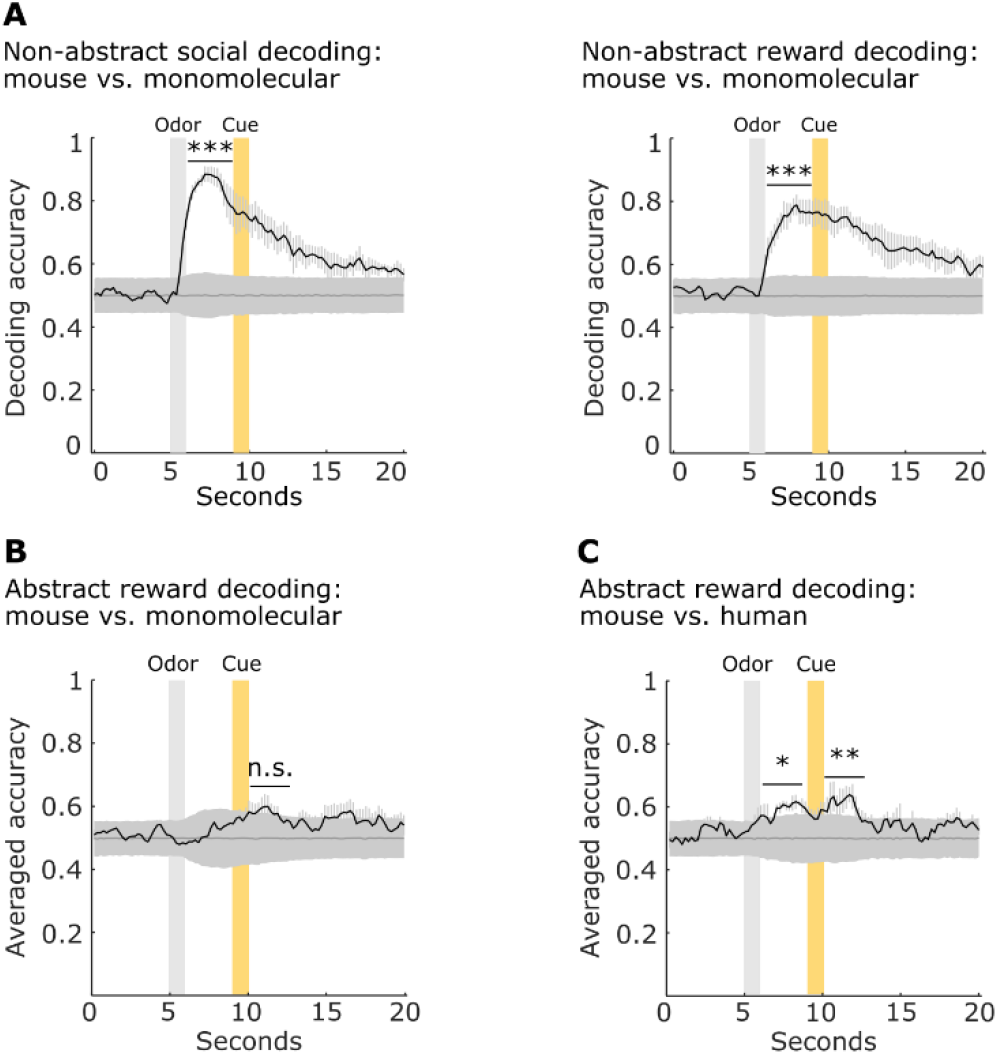
Non-abstract decoding for mouse versus monomolecular odors and abstract reward decoding. (A) Left, average decoding accuracy of a SVM binary linear decoder trained and tested on all two rewarded and two non-rewarded odors to distinguish social from non-social category (70 cells, 1,000 samples per mouse; n = 5). Significance is determined in the three second delay window after odor presentation by calculating an empirical p-value using the pairwise difference distribution between signal and chance distributions; right, same as left, but the decoder was trained to distinguish rewarded from non-rewarded odors. ***, p < 0.001. (B) Decoding accuracy across the reward/non-reward category dimension as a function of time. The black trace shows the mean decoding accuracy of the two linear decoders (SVM) as outlined in (F), averaged over three mice; mean ± SEM. The first decoder was trained to distinguish between rewarded and non-rewarded social odors (urine G and urine H). The trained model was then asked to classify the two distinct odors of the remaining pair: rewarded versus non-rewarded non-social odors (methyl butyrate and ethyl acetate). The second decoder was trained to distinguish rewarded from non-rewarded non-social odors and was then tested on the pair of rewarded odors. Chance decoding was estimated by decoding with shuffled trial information; n.s., non-significant. (C) Same as in (B) except using a novel set of mouse urine I and J and two human urine samples as non-social odors; n = 3, **, p < 0.01, *, p < 0.05.

